# Cell Surface-localized CsgF Condensate is a Gatekeeper in Bacterial Curli Subunit Secretion

**DOI:** 10.1101/2022.10.20.513119

**Authors:** Hema M. Swasthi, Joseph L. Basalla, Claire E. Dudley, Anthony G. Vecchiarelli, Matthew R. Chapman

## Abstract

Curli are functional amyloids present on the outer membrane of *E. coli*. Cell-surface aggregation of CsgA, the major subunit of curli, is a well-orchestrated process. CsgB, the minor subunit of curli, nucleates the aggregation of CsgA while CsgF, a curli accessory protein, ensures proper anchoring of CsgB to the cell surface. The molecular basis of the interactions between CsgF and curli subunits is unclear. Here, we show that CsgF undergoes phase separation *in vitro* and that the ability of CsgF variants to phase separate tightly correlated with CsgF function in cells during curli biogenesis. Substitution of phenylalanine residues in the CsgF N-terminus both reduced the propensity of CsgF to phase-separate and impaired curli biogenesis. Exogenous addition of purified CsgF complemented *csgF* ^−^ cells. This exogenous addition assay was used to assess the ability of CsgF variants to complement *csgF* ^−^ cells. The presence of CsgF on the cell surface modulated the secretion of CsgA to the cell surface. We also found that the CsgB nucleator protein is a CsgF client. CsgB can form SDS-insoluble aggregates within the dynamic CsgF condensate, and we propose that these multi-component CsgF-B condensates form a nucleation-competent complex that templates CsgA amyloid formation on the cell surface. Together, our study provides insight into the ability of CsgF to phase separate, regulate CsgA secretion, and promote CsgB aggregation in curli assembly.

## Introduction

Protein misfolding and amyloid deposition are implicated in many neurodegenerative disorders such as Parkinson’s disease and Alzheimer’s disease ^1,2^. ‘Functional’ amyloids are distinguished in that the amyloid structure is tightly linked to a desired physiological outcome for the cell ^3–5^. Curli fibers are functional amyloids found in the extracellular matrix of enteric bacteria such as *E*.*coli* and *Salmonella* ^6^. Curli are involved in biofilm formation, cell-cell adhesion, cell-host interaction, and can trigger immune responses in host cells ^6,7^. Curli amyloids are composed of a major subunit called CsgA (<u>c</u>urli <u>s</u>pecific <u>g</u>ene) and a minor subunit called CsgB ^8^. CsgA amyloid formation at the cell surface is initiated by the CsgB ‘nucleator’ protein. ^8–10^. Five accessory proteins are necessary to regulate the secretion of curli subunits and to ensure the proper formation of curli on the cell surface. The curli subunits and the accessory proteins are divergently transcribed from the operons *csgDEGF* and *csgBAC* ^11–13^. CsgD, the curli master regulator, regulates the expression of all curli-associated proteins ^14^. CsgG is a nonameric membrane pore that spans the outer membrane and serves as the channel for the secretion of CsgA, CsgB, and CsgF ^15–17^. The binding of CsgE to the periplasmic region of CsgG is essential for substrate selection ^16,18^. CsgC is a chaperone-like protein that prevents the aggregation of CsgA in the periplasmic region of the bacterial cell ^19^. CsgA is secreted to the outer membrane in a soluble unpolymerized form ^20^. On the cell surface, CsgB nucleates CsgA amyloid formation ^8^. The accessory protein CsgF is essential for the proper anchoring of CsgB onto the cell surface, and without CsgF curli formation is disrupted ^12,21^. Recent structural studies using cryo-EM have revealed that the N-terminus of CsgF arranges on the extracellular part of CsgG ^22–24^. However, the mechanism by which CsgF aids the curli assembly is poorly understood.

The classical view on bacterial cell organization is rapidly evolving ^25^. A growing body of evidence suggests that biomolecules can localize within bacterial cells ^25–27^. Membraneless sub-compartmentalization of biomolecules into “biocondensates” via phase separation (density transition) or phase separation coupled to percolation (networking transition) has emerged as a common theme in eukaryotic cells ^28–34^. Advances in microscopy techniques have revealed that the intracellular organization of biomolecules in bacteria could also occur via phase separation ^27,35,36^. Multivalent weak interactions between proteins or protein-nucleic acids facilitate the assembly of biocondensates ^30,37,38^. Proteins with low-complexity domains, intrinsically disordered regions (IDRs), or prion-like regions are shown to undergo phase separation however, the role of these domains/regions to mediate phase separation depends on its amino acid composition ^39–41^. Biocondensates have properties that are distinct from their surroundings and have been shown to be involved in a wide range of cellular functions ^28,42^. For example, concentrating specific reactants in biocondensates can enhance biochemical reactions or suppress reactions by sequestering reactants ^37,43^. Many neurodegenerative disease-associated amyloid forming proteins, such as α-synuclein, huntingtin, tau, and the prion proteins, have been shown to phase-separate and aggregate into amyloids within biocondensates ^20,28,30,44–46^.

The bacterial functional amyloid curli shares biochemical and biophysical properties with diseases-associated amyloids ^6^. Cell-surface aggregation of CsgA, the major structural component of curli, is nucleated by CsgB ^9^. CsgB adopts a protease-resistant structure on the cell surface ^21^. CsgF is the curli accessory protein secreted to the outer membrane that helps CsgB to anchor on the cell surface and to attain nucleator activity ^21^. The mechanism by which CsgF promotes the nucleator activity of CsgB is not well understood. In this study, we demonstrate that CsgF readily phase separates *in vitro* and that aromatic residues in the N-terminal region of CsgF are critical for both its phase separation activity *in vitro* and curli biogenesis *in vivo*. We find that CsgF on the cell surface regulates the amount of CsgA that is secreted to the outer membrane. Additionally, we show that the curli nucleator protein CsgB associates with CsgF droplets and undergoes an amyloid transition while CsgF remains dynamic. Taken together, this study sheds light into the interplay between CsgF and curli subunits during the formation of functional amyloids on bacteria.

## Results

### CsgF forms biocondensates *in vitro*

Mature CsgF has 119 amino acids, with the first 19 amino acids of CsgF processed during translocation to the periplasm (Figure 1A). NMR studies on CsgF have shown that both the N-terminus (CsgF_N_) and C-terminus (CsgF_C_) are unstructured, while the middle region (CsgF_M_) is a mixture of α-helices and β-strands (Figure 1A, 1B, and S1A) ^47^. CsgF is rich in asparagine (16 residues) and glutamine (11 residues) residues, which is one of the characteristics of prion proteins ^48^. Therefore, we subjected CsgF to prion-like amino acid composition (PLAAC) analysis ^49^. PLAAC predicted that the N-terminus of CsgF (19 to 54) is prion-like (Figure 1C). Multivalent interactions mediated by low-complexity <u>p</u>rion-like <u>d</u>omains (PLDs) have been shown to drive phase separation of proteins ^39,50^. Therefore, we asked whether CsgF can undergo phase separation. Purified wild-type (WT) CsgF was incubated at room temperate at pH 7.5 before turbidity was assessed by eye (Figure 1D), or at 350 nm using a spectrometer (Figure 1E). Turbidity increased as the WT-CsgF concentration increased from 2.5 μM to 100 μM (Figure 1E). Next, lysine residues of WT-CsgF were sparsely labeled with Alexa-633 and mixed with unlabeled protein to a 1:50 (labeled:unlabeled) molar ratio. Fluorescence imaging revealed the formation of phase-separated droplets of WT-CsgF in a concentration-dependent manner (Figure 1F). The minimum concentration of WT-CsgF required to undergo measurable phase separation was 5 μM (Figure 1F). WT-CsgF droplets fused to form larger droplets (Movie 1). Fluorescence recovery after photobleaching (FRAP) measurements on WT-CsgF condensates revealed ∼ 60% recovery with t_1/2_ = 15 ± 4 s (Figure 1G). Interestingly, fluorescence recovery of 20 hr old (t_1/2_ = 47 ± 14 s) WT-CsgF droplets was comparable to that of 1 hr old WT-CsgF droplets (t_1/2_ = 36 ± 8 s) (Figure 1G), suggesting that WT-CsgF condensates do not mature into a more viscous state over time.

**Figure 1.**
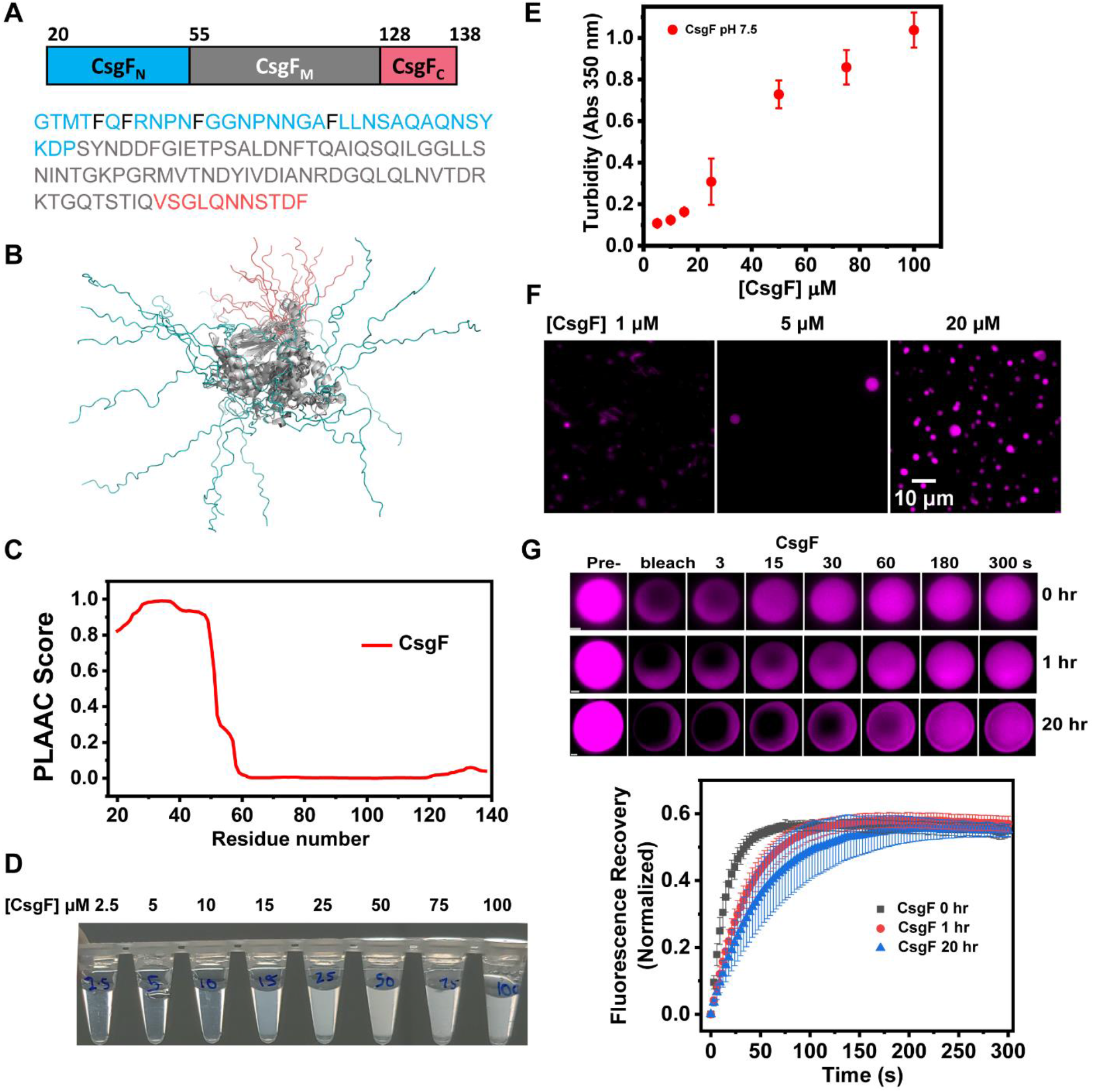
CsgF undergoes phase separation. **(A)** Schematic representation and amino acid sequence of mature CsgF. The N-terminal region (amino acids 20-54) is colored in blue, the middle domain (amino acids 55-127) is shown in grey, and the C-terminal region (amino acids 128-138) is indicated in pink. Phenylalanine residues in the N-terminus are indicated in black. **(B)** An overlay of 16 conformations of CsgF NMR structure (PDB code: 5M1U). **(C)** PLAAC analysis on CsgF. **(D)** Concentration-dependent density change of CsgF in 25 mM potassium phosphate pH 7.5 buffer. **(E)** Concentration-dependent turbidity change of CsgF, 25 mM potassium phosphate pH 7.5 measured at 350 nm. **(F)** Fluorescence images of CsgF at 1 μM, 5 μM, and 20 μM in 25 mM potassium phosphate pH 7.5. The unlabeled to labeled CsgF ratio was 50:1. **(G)** Time-dependent FRAP measurements on CsgF and quantification of FRAP recovery as a function of time. The error bars shown are the standard deviation (SD) of FRAP measurements obtained from 10 or more droplets.

We next investigated how pH affected the propensity of WT-CsgF to undergo phase separation. The turbidity of WT-CsgF was measured between a pH range of 4.0 to 8.5. Below pH 4.5 and above pH 8 there was no apparent turbidity changes, even at the highest concentrations of WT-CsgF (Figure S1B). The pI of WT-CsgF is 4.8 and the predicted net charge of WT-CsgF at pH 4 and 8.5 is +4.3 and -3.9, respectively. We speculated that charge repulsion could prevent WT-CsgF from phase separating at pH 4 and at 8.5. Salt can mask the surface charge of proteins, therefore sodium chloride was added to WT-CsgF at pH 4 to access changes in its phase separation activity. The addition of 200 mM NaCl induced phase separation of WT-CsgF at pH 4 (Figure S1C). The turbidity of WT-CsgF at pH 4 in the presence of 250 mM NaCl increased as the protein concentration was increased (Figure S1D). At 400 mM NaCl, WT-CsgF formed solid-like aggregates (Figure S1C). Thus, addition of salt to WT-CsgF at pH 4 could be weakening the ionic interactions and enhancing the non-ionic interactions, which results in phase separation or aggregation of WT-CsgF ^51^. Collectively, this data suggested that shielding the surface charge of WT-CsgF promoted its phase separation, which implicates non-ionic interactions as important for WT-CsgF phase separation.

### Phenylalanine residues in the N-terminus modulate CsgF phase separation

Structural elucidation of WT-CsgF using NMR has shown that both the N- and C-termini are unstructured (Figure 1B) ^47^. We made N- and C-terminal truncation mutants of CsgF to determine if either IDR was necessary for its phase separation activity (Figure 2A). The turbidity of CsgF-ΔC samples increased as the protein concentration increased (Figure 2B), and imaging confirmed condensate formation (Figure 2C). CsgF-ΔN, on the other hand, did not show concentration-dependent turbidity changes (Figure 2B) or visible droplet formation, even at significantly higher protein concentrations (300 μM) (Figure 2C). WT-CsgF and its variants (20 μM and 50 μM) were centrifuged and the supernatant and pellet fractions were subjected to SDS-PAGE to quantify proteins in the dilute and condensed phases (Figure S2A and 2D). The amount of CsgF-ΔC in the pellet fraction was reduced by ∼ 20% relative to WT-CsgF, while most of the CsgF-ΔN remained in the supernatant (Figure 2E). Therefore, both the N- and C-terminal disordered regions of CsgF contribute to its phase separation activity. However, the N-terminus appears to be necessary for the phase separation activity of WT-CsgF, while the C-terminus is dispensable, at least at relatively high protein concentrations. The phase separation of CsgF-ΔN was monitored in the presence of a crowding agent to elucidate whether WT-CsgF can phase separate without the disordered N-terminus. Phase separation was not observed upon addition of 5% PEG-8000 to 100 μM CsgF-ΔN (Figure 2F). However, 5% crowding agent induced phase separation of 300 μM CsgF-ΔN (Figure 2F), which suggested that WT-CsgF can phase separate even in the absence of the N-terminal IDR. Thus, increasing the local concentration of CsgF-ΔN with a crowding agent promoted its phase separation. However, WT-CsgF phase separated at a concentration of ∼ 5 μM while CsgF-ΔN required an approximately 60-fold increase in the protein concentration and the addition of PEG-800 to phase separate. The N-terminal region might be bringing WT-CsgF together to increase the effective protein concentration and could also be involved in weak multivalent interactions.

**Figure 2.**
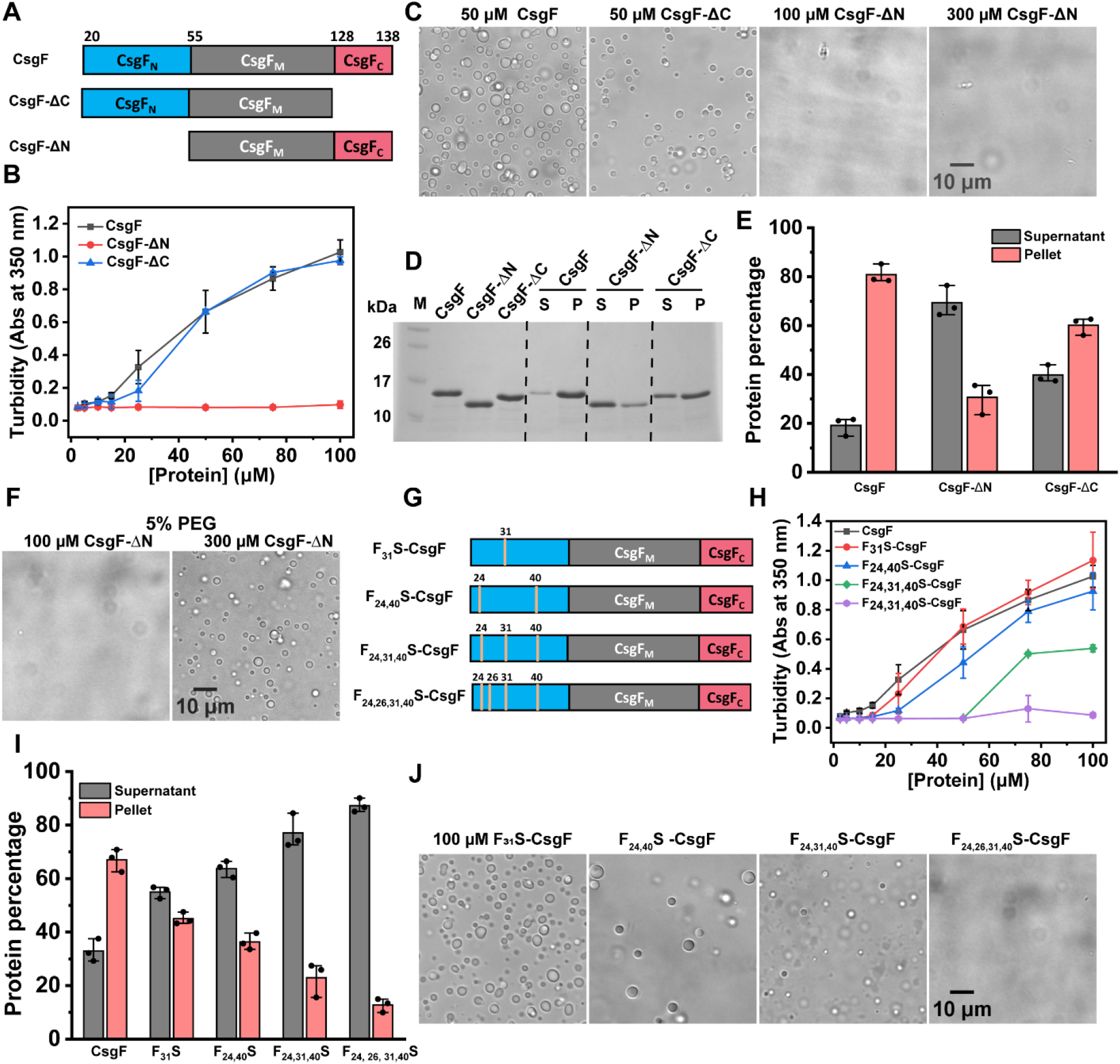
Phenylalanine residues in the N-terminus modulate phase separation of CsgF. **(A)** Schematic representation of CsgF-ΔN and CsgF-ΔC used in the study. **(B)** Turbidity measurements of CsgF, CsgF-ΔN, and CsgF-ΔC in 25 mM potassium phosphate pH 7.5 **(C)** Images of CsgF (50 μM), CsgF-ΔC (50 μM) and CsgF-ΔN (100 μM and 300 μM) in 25 mM potassium phosphate pH 7.5. **(D)** Coomassie-stained SDS-PAGE on the supernatant and pellet samples of 50 μM CsgF, CsgF-ΔN, and CsgF-ΔC after centrifugation. **(E)** SDS-PAGE shown in 4D was quantified. **(F)** DIC images of 100 μM and 300 μM CsgF-ΔN in the presence of 5% PEG-8000. **(G)** The phenylalanine residues that are mutated in the N-terminal region are marked as yellow lines. **(H)** Turbidity measurements on phenylalanine mutants of CsgF. **(I)** Sedimentation assay on the phenylalanine mutants of CsgF was quantified from Coomassie-stained SDS-PAGE. The error bars are the SD obtained from three independent measurements. **(J)** DIC images of 100 μM CsgF phenylalanine mutants in 25 mM potassium phosphate pH 7.5.

Studies have demonstrated that aromatic residues in IDRs determine the phase behavior of proteins ^52–54^. WT-CsgF has a total of 7 phenylalanine and 3 tyrosine residues, with four phenylalanine residues and one tyrosine residue in the N-terminal region (Figure 1A). We created serine substitutions of the phenylalanine residues located in the CsgF N-terminus. Phenylalanine to serine mutants were named F_X_S-CsgF (X= the position where the original phenylalanine was present) (Figure 2G). To measure how the phase separation activity of WT-CsgF varies with changes to the number of phenylalanine residues, we first used a turbidity assay. Turbidity measurements indicated that as the number of phenylalanine residues decreased there was a corresponding reduction in WT-CsgF phase separation (Figure 2H). F_24,26,31,40_S-CsgF did not show any measurable turbidity (Figure 2H). Consistently, sedimentation assays performed on 20 μM F_X_S-CsgF mutants revealed a progressive reduction in the amount of protein in the pellet fraction with the decrease of phenylalanine residues (Figure S2B and 2I). We imaged the phenylalanine to serine mutants at 50 μM and 100 μM concentrations and found that F_31_S and F_24,40_S-CsgF readily phase separated at these concentrations (Figure S2C and Figure 2J). F_24,31,40_S-CsgF, on the other hand, only phase separated at 100 μM. F_24,26,31,40_S-CsgF showed no phase separation activity at either of the protein concentrations tested (Figure S2C and 2I). The sedimentation assays, turbidity measurements, and microscopic imaging suggested that the phenylalanine residues in the N-terminus are required to promote phase separation of WT-CsgF.

### The N-terminus is essential for the secretion of CsgF to the outer membrane

CsgF assists CsgB to form a functional curli nucleator complex on the bacterial surface ^21^. In the absence of CsgF, cells fail to assemble as much curli on the cell surface as wild-type (WT) ^12,21^. We set out to understand the role of the N- and C-terminal IDRs of CsgF by assaying the ability of CsgF-ΔN and CsgF-ΔC variants to complement WT-CsgF function *in vivo*. Curli producing *E*.*coli* cells MC4100 (WT/empty vector (EV)) stain red on Congo red (CR) indicator plates, and the *csgF* ^*-*^/EV cells exhibit a pink color phenotype (Figure 3A) ^12,21^. When *csgF* ^*-*^/EV cells were removed from CR plates after 48 hours of growth, the underlying agar was red (Figure 3A) ^21^. The red coloration underneath the *csgF* ^*-*^*/*EV cells is due to secreted curli subunits that assemble into CR-binding polymers, but did not attach to the cells ^21^. The *csgF* ^*-*^ strain was complemented by the pCsgF plasmid (Figure 3A). However, when *csgF* ^*-*^ cells harboring pCsgF-ΔN were grown under curli expressing conditions the phenotype matched the *csgF* ^*-*^ strain where the underlying agar was stained red. Interestingly, the *csgF* ^*-*^/ pCsgF-ΔC cells remained white even after 48 hr, and when the cells were scraped off there was no red staining on the agar (Figure 3A). Thus, failure of CsgF-ΔN and CsgF-ΔC to complement *csgF* ^*-*^ cells indicated that both N- and C-terminal regions of CsgF are necessary for the function of WT-CsgF.

**Figure 3.**
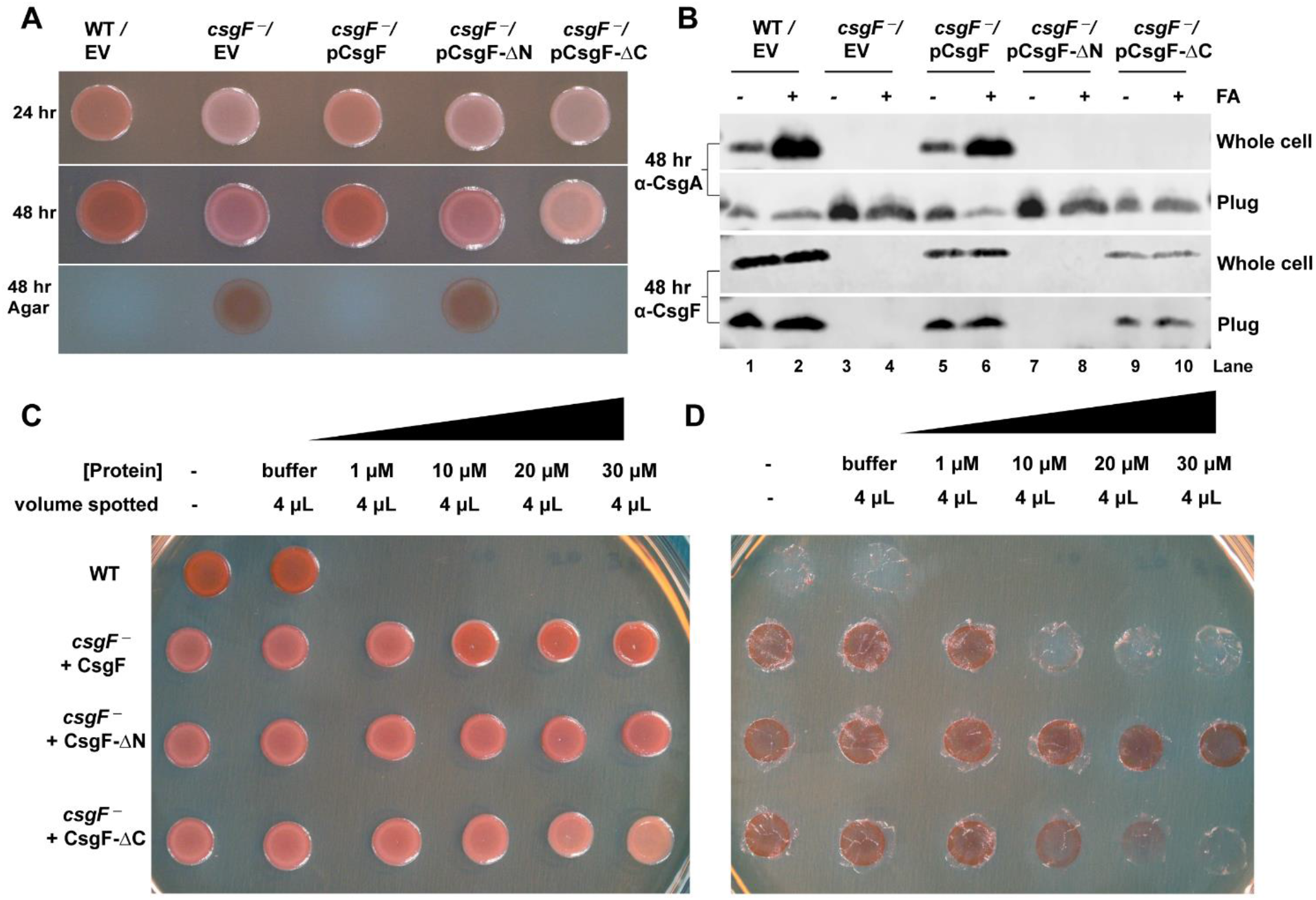
Exogenous CsgF complements *csgF* ^*-*^ cells. **(A)** Cells were grown on Congo red (CR) indicator plate to assess the phenotypes of WT/Empty vector (MC4100/pLR1), *csgF* ^*-*^/Empty vector (MHR592/pLR1), *csgF* ^*-*^/pCsgF (MHR592/pLR73), *csgF* ^*-*^/pCsgF-ΔN (MHR592/pCsgF-ΔN), and *csgF* ^*-*^/pCsgF-ΔC (MHR592/pCsgF-ΔC), and after 24 hr and 48 hr and agar plate after removing bacterial cells. **(B)** Western blots on whole cells and plugs after 48 hr of growth on YESCA plates. The samples for blots were prepared either in 2X SDS-PAGE running buffer directly, or first being treated with formic acid (FA) before being run on SDS-PAGE, transferred to membrane, and probed for CsgA and CsgF. **(C)** *csgF* ^*-*^ cells complemented using exogenously added protein. Protein stocks (1 μM, 10 μM, 20 μM, and 30 μM) were made in buffer (25 mM Tris pH 9). Four μL (1 μM, 10 μM, 20 μM, and 30 μM) of protein was spotted on the Congo red YESCA plate and when the protein sample was dried, cells were spotted and allowed to grow for 48 hr at 26 ºC. **(D)** Agar plate after removing the bacterial cells shown in 3C.

To better understand how the N- and C-terminal domains contribute to WT-CsgF function *in vivo, csgF* ^*-*^ strains harboring pCsgF-ΔN and pCsgF-ΔC were characterized. Curli-producing WT cells and *csgF* ^*-*^/pCsgF cells need mild formic acid treatment to monomerize the cell-surface polymerized CsgA ^12,21^. CsgA that is secreted by curli-forming cells is assembled as amyloid fibers on the outer membrane, thus, there would not be much SDS-soluble CsgA in the agar plug. In *csgF* ^*-*^ strains, CsgA fails to polymerize on the cell surface allowing unpolymerized CsgA to secrete away from the cell surface ^21^. Therefore, the agar-plug underneath *csgF* ^*-*^/EV cells have SDS-soluble CsgA. Western blot analysis on whole cells and the plug was performed to characterize the nature of CsgA in cells that were expressing CsgF variants. Without formic acid treatment, very little monomeric CsgA was detectable in the WT/EV or *csgF* ^*-*^/pCsgF cells that were collected from YESCA plates after 48 hours of growth (Figure 3B, top). Upon treating the WT/EV and *csgF*^*-*^/pCsgF cells with formic acid there was a dramatic increase in the amount of CsgA in the whole cell blot suggesting that CsgA was in an SDS-insoluble amyloid form (Figure 3B). On the contrary, for *csgF* ^*-*^/EV cells, CsgA was found in the plugs, and formic acid treated and non-treated plugs had a comparable amount of CsgA (Figure 3B). Similarly, *csgF* ^*-*^ cells with pCsgF-ΔN and pCsgF-ΔC, CsgA was neither detected in formic acid-treated nor in non-treated whole cells. Both in pCsgF-ΔN and pCsgF-ΔC containing strains, CsgA was found in the agar plugs. We next probed for CsgF (Figure 3B, bottom). WT/EV and *csgF* ^*-*^/pCsgF cells showed the presence of WT-CsgF both in the whole cells and plug. However, *csgF* ^*-*^/pCsgF-ΔN did not have detectable bands in either the whole cell or plug samples when stained with CsgF antibody (Figure 3B, lanes 7 and 8). *csgF* ^*-*^ cells with pCsgF-ΔC were detected by CsgF antibody both in the whole cell and plug samples. To test whether CsgF antibody binds to CsgF-ΔN, we probed the antibody against purified proteins. CsgF antibody staining of CsgF-ΔN matched with WT-CsgF but CsgF antibody exhibited slightly less binding to CsgF-ΔC (Figure S3A). Thus, the absence of CsgF-ΔN in the whole cell or plug was not because of the inability of CsgF antibody to detect CsgF-ΔN variants as the CsgF antibody exhibited similar affinity towards WT-CsgF and CsgF-ΔN (Figure S3A). The absence of CsgF-ΔN both in the whole cell and plug suggested that CsgF-ΔN is unstable and not secreted to the cell surface.

### Phenylalanine residues are critical for curli assembly

Since CsgF is a cell surface protein we hypothesized that exogenous WT-CsgF would complement *csgF* ^*-*^ cells. To test this, *csgF* ^*-*^ cells were spotted on a CR-YESCA plate that had been supplemented with purified WT-CsgF. After 48 hours of growth, the *csgF* ^*-*^ cells appeared pink, but as the concentration of purified WT-CsgF that was added to the plate increased, the cells stained a deeper red (Figure 3C). Similarly, there was a decrease in the red staining on the CR plate as the concentration of exogenously added WT-CsgF increased (Figure 3D). Thus, exogenous WT-CsgF partially complemented *csgF* ^*-*^ cells. When CsgF-ΔN protein was added to the *csgF* ^*-*^ cells, the phenotype was similar to *csgF* ^*-*^ cells even at the highest protein concentration tested (1.12 μg). *csgF* ^*-*^ cells grown in the presence of 1.54 μg of CsgF-ΔC variant exhibited a white phenotype and the red coloration on the plate decreased with an increase in protein concentration (Figure 3C and D). Thus, the phenotypic features exhibited by *csgF* ^*-*^/pCsgF, *csgF* ^*-*^/pCsgF-ΔN, and *csgF* ^*-*^/pCsgF-ΔC were recapitulated using endogenous WT-CsgF or its variants. The absence of CsgF-ΔN in the whole cell and plug, as well as the phenotypic similarity of *csgF* ^*-*^ /pCsgF-ΔN and *csgF* ^*-*^ suggested that the N-terminal region of CsgF is required for the function of protein on the cell surface. The failure of endogenous and exogenous CsgF-ΔN and CsgF-ΔC to complement *csgF* ^*-*^ cells indicated that both the N-terminal and C-terminal regions of CsgF are essential for the function of the protein. The N-terminus of CsgF has a key role to play both during CsgF translocation across the outer membrane and on the cell surface during curli biogenesis.

Next, we focused on elucidating the ability of phenylalanine mutants (Figure 2G) to complement *csgF* ^*-*^ cells. We used the exogenous complementation method to assess the complementation ability of the phenylalanine mutants. None of the phenylalanine mutants complemented *csgF* ^*-*^ cells (Figure 4A). Intriguingly, a single F31S mutation was enough to impair the function of WT-CsgF and curli assembly. We also examined single phenylalanine mutation at 24, 26, and 40 positions. CR assay revealed that all the single phenylalanine mutations failed to complement *csgF* ^*-*^ cells (Figure 4B). Phase separation activity was assessed to see whether single phenylalanine mutation affected the phase separation propensity of WT-CsgF. 20 μM samples were sedimented by centrifugation and analyzed on a SDS-PAGE (Figure 4C and D). There was a reduction in the amount of protein present in the pellet fraction of all single F_x_S mutants compared to the WT-CsgF. Imaging of the single phenylalanine mutants suggested all the single phenylalanine mutants phase separated ∼ 20 μM whereas, WT-CsgF phase separates ∼ 5 μM (Figure 4E and S3B). The CR plate assay, together with sedimentation and imaging, suggested that the phenylalanine residues in the CsgF N-terminus are critical for WT-CsgF function and phase separation.

**Figure 4.**
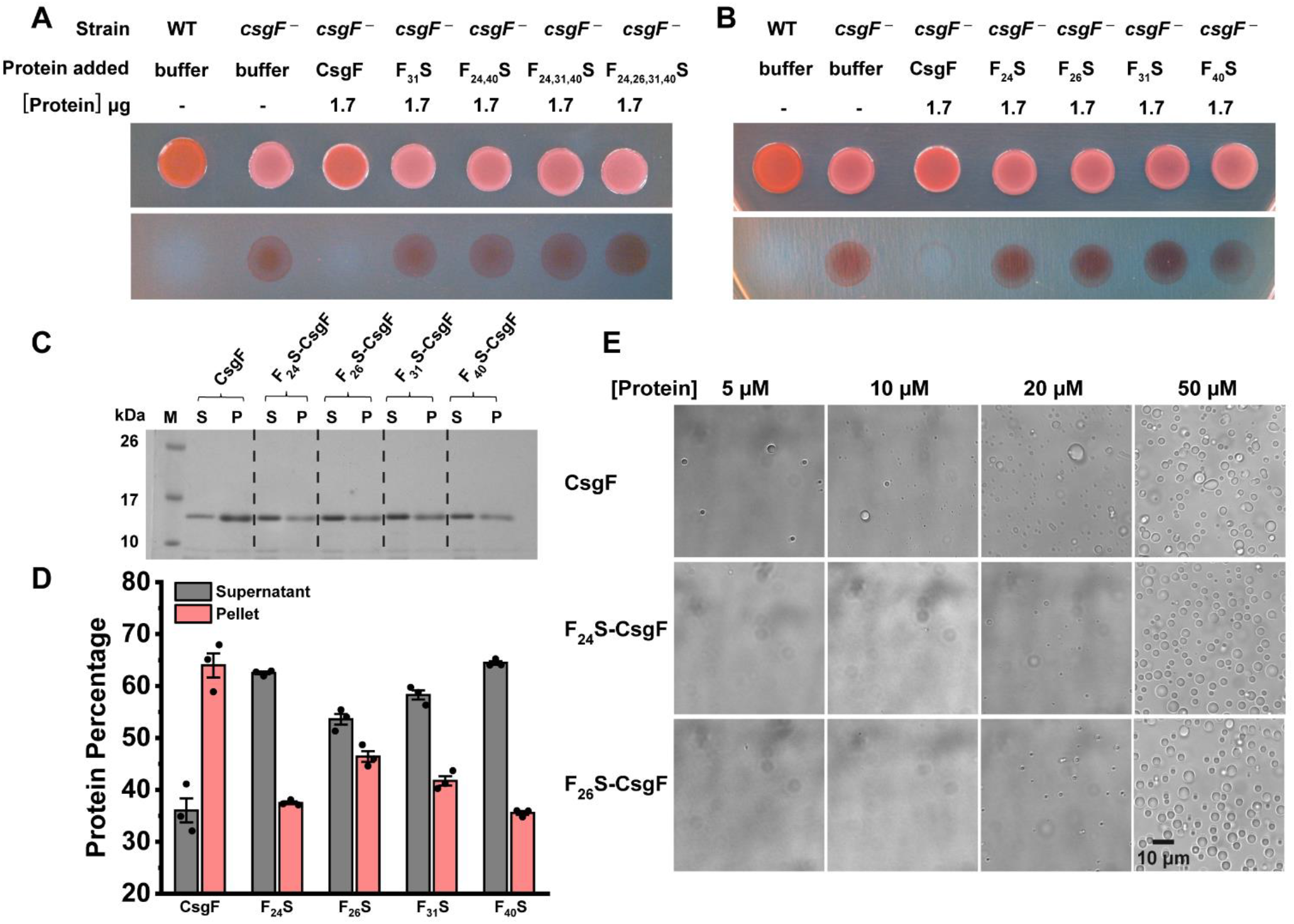
Single phenylalanine mutation impairs curli assembly and CsgF phase separation. **(A)** and **(B)** Exogenous complementation of *csgF* ^*-*^ cells with phenylalanine mutants. **(C)** Coomassie-stained SDS-PAGE on the supernatant and pellet samples of single phenylalanine mutants after sedimentation. **(D)** Quantification of SDS-PAGE shown in 4C. **(E)** DIC images of 5 μM, 10 μM, 20 μM, and 50 μM of CsgF, F_24_S-CSgF, and F_26_S-CsgF in 25 mM potassium phosphate pH 7.5.

### Surface-localized CsgF regulates CsgA secretion

The unpolymerized soluble CsgA that is secreted away from a *csgB* ^*-*^ strain (CsgA^+^ donor) can assemble on the surface of *csgF* ^*-*^ strain (CsgB^+^ recipient) by the process of interbacterial complementation ^55^. The *csgFB* ^*-*^ double mutant cells are better CsgA donors compared to *csgB* ^*-*^ cells ^12^ and why *csgFB* ^*-*^ cells are better CsgA donors is unknown. We performed a CR assay on CsgA donors *csgFB* ^*-*^/EV, *csgFB* ^*-*^*/*pCsgF, *csgB* ^*-*^/EV, along with *csgF* ^*-*^ and its variants. The *csgF*^*-*^/EV and *csgF* ^*-*^/pCsgF-ΔN cells were pink on CR indicator plates, while all the other strains remained white or unstained (Figure 5A and 5B). Only *csgF* ^*-*^/EV and *csgF* ^*-*^/pCsgF-ΔN stained red underneath the cells. To see the presence of CsgF on the bacterial surface we carried out intact cell dot blot assays. Cells grown on YESCA plates for 48 hr were normalized to 1 OD_600_ and spotted on nitrocellulose membrane and probed for CsgF. As expected, WT/EV, *csgFB* ^*-*^/pCsgF, *csgB* ^*-*^/EV, *csgB* ^*-*^/pCsgB, *csgF* ^*-*^/pCsgF, and *csgF* ^*-*^/pCsgF-ΔC showed the presence of surface-associated CsgF (Figure 5C). Next, we performed a plug assay without formic acid to determine the amount of soluble CsgA that was being secreted by these non-curli producing cells. As a loading control, we ran the samples shown in figure 5D on SDS-PAGE and stained with Coomassie blue (Figure S3C). Interestingly, we observed differences in the amount of CsgA in the plug for the non-curli producing cells (Figure 5D and 5E). The amount of CsgA secreted by the *csgFB* ^*-*^/EV double mutant cells was higher compared to other non-curli producing cells. *csgB* ^*-*^/EV cells secreted ∼ 2.5-fold less CsgA relative to the *csgFB* ^*-*^/EV cells (Figure 5E). Similarly, *csgF* ^*-*^/pCsgF-ΔC secreted less CsgA protein than *csgFB* ^*-*^/EV cells. Thus, the presence of cell surface CsgF in the non-curli producing cells governed the CsgA secretion. Next, we performed immunofluorescence microscopy to visualize cell surface associated CsgF. Immunofluorescence imaging of *csgF* ^*-*^/pCsgF-His revealed that WT-CsgF localized as puncta on the cell surface (Figure 5F). The plug and immunofluorescence assays, together with the intact cell dot blot, indicated that WT-CsgF localizes to the bacterial surface and is a regulator of CsgA secretion in the non-curli producing cells (Figure 5G).

**Figure 5.**
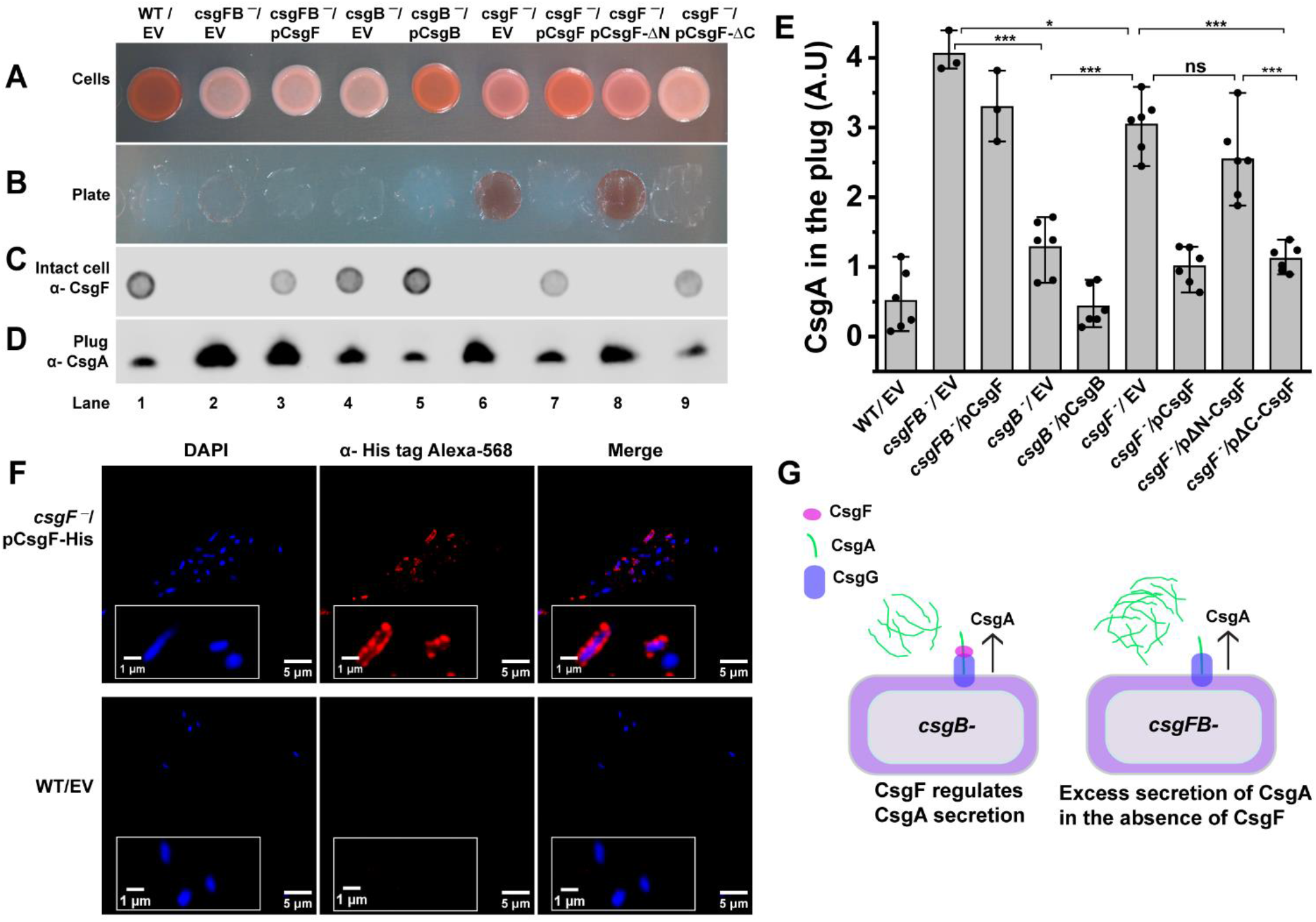
Cell surface-associated CsgF regulates CsgA secretion. **(A)** Phenotypes of 48 hr grown WT/Empty vector (MC4100/pLR1), *csgFB* ^*-*^/Empty vector (*csgFB* ^*-*^/pLR2), *csgFB* ^*-*^/pCsgF (*csgFB* ^*-*^/pLR73), *csgB* ^*-*^/Empty vector (MHR261/pLR2) *csgB* ^*-*^/pCsgB (MHR261/pLR8), *csgF* ^*-*^/Empty vector (MHR592/pLR1), *csgF* ^*-*^/pCsgF (MHR592/pLR73), *csgF* ^*-*^/pCsgF-ΔN (MHR592/pCsgF-ΔN), and *csgF* ^*-*^/pCsgF-ΔC (MHR592/pCsgF-ΔC) cells on a Congo red YESCA plate. **(B)** Agar plate after removing the cells shown in (A). **(C)** Dot blot was performed using CsgF antibody on intact cells to assess the presence of surface-associated CsgF. Four microliters of a 1 OD_600_ stock of cells were spotted on the nitrocellulose membrane and probed with CsgF antibody. **(D)** Western blot on agar plugs that were collected from under the cells after they had grown for 48 hr and removed. **(E)** Western blot shown in D was quantified using image J. Significance was determined by one-way ANOVA with the Tukey test. * p<0.05, ***p<0.001 **(F)** Immunofluorescence images of *csgF* ^*-*^/pCsgF-His (MHR592/pCsgF-His). The cells were fixed with formaldehyde and probed with His-tag antibody and then with Alexa-568 secondary antibody. WT/Empty vector (MC4100/pTrc99A) was used as a control. **(G)** A model to show the presence of surface-associated CsgF regulates CsgA secretion.

### CsgB incorporates into phase-separated droplets of CsgF

Curli subunits are secreted to the cell surface via the CsgG membrane channel. Cryo-EM structural analysis has shown that the N-terminal region of CsgF binds to the CsgG channel ^22–24^. CsgB and CsgA could potentially interact with the CsgG-CsgF complex while traversing to the cell surface. In the absence of WT-CsgF, CsgB fails to anchor on the cell surface and nucleate CsgA polymerization ^21^. We asked whether CsgB could interact with the WT-CsgF condensates we identified here. When CsgF (20 μM) condensates were added to CsgB (20 μM), both proteins colocalized (Figure 6A). However, CsgB alone did not undergo phase separation under WT-CsgF condensate-forming conditions (Figure S4A). The C-terminal region (R5 repeat) of CsgB is positively charged and a CsgB truncation mutant that is missing R5 (CsgBΔR5) does not aggregate into amyloid fibers as readily as WT-CsgB ^10,56^. CsgBΔR5 still sequestered into WT-CsgF phase-separated droplets (Figure S4B). CsgB shares ∼ 30% sequence identity with CsgA ^9^. To see whether the interaction between CsgB and WT-CsgF was specific, we also performed fluorescence imaging of WT-CsgF condensates in the presence of CsgA. Unlike CsgB, CsgA was not recruited into CsgF condensates (Figure S4C). To elucidate the nature of CsgB in the phase-separated droplets of WT-CsgF, we carried out FRAP measurements. CsgB recruited in the WT-CsgF droplets exhibited a slow and linear fluorescence recovery immediately after mixing (Figure 6B and 6C). The half time recovered from the fluorescence recovery kinetics of CsgF immediately after mixing with CsgB and after an hour were, 32 ± 6 s, and 56 ± 15 s, respectively (Figure 6D and 6E). CsgB, on the other hand, failed to show any notable fluorescence recovery after an hour. SDS-PAGE was performed to further understand the nature of CsgB in the presence of WT-CsgF. Samples collected after 10 min contained SDS-soluble CsgB (Figure 6F). However, after 30 and 60 min the samples had very little soluble CsgB and most of the CsgB was stuck in the SDS-PAGE wells (Figure 6F). The SDS-PAGE data suggested the formation of SDS insoluble higher order CsgB aggregates in the presence of phase separated CsgF droplets. It is noteworthy that the material state of CsgB selectively transitioned from a dynamic to an amyloid state in the multicomponent droplets while, WT-CsgF did not undergo notable changes. Fluorescent imaging of curli subunits with CsgF indicated that the interaction between CsgB and WT-CsgF droplets are specific and that CsgB undergoes amyloid transition within the dynamic CsgF condensates.

**Figure 6.**
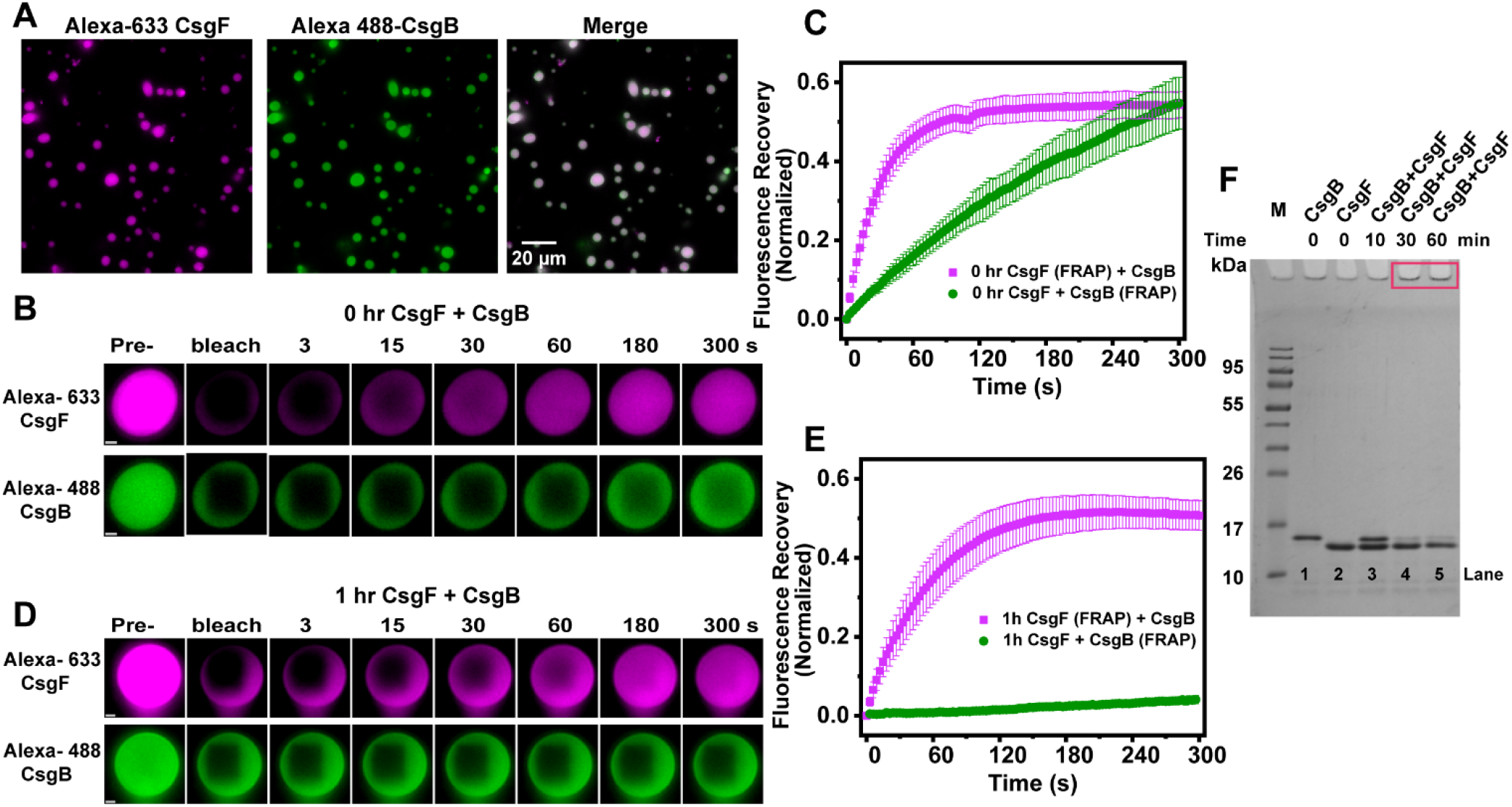
CsgB localizes in CsgF condensates. **(A)** Fluorescence images of Alexa-633-CsgF (20 μM) and Alexa-488-CsgB (20 μM). Unlabeled to labeled protein molar ratio used was 50:1. **(B)** FRAP on CsgF and CsgB immediately after mixing. **(C)** Quantification of fluorescence recovery of CsgF and CsgB for 0 hr sample. (D) FRAP measurements on CsgF and CsgB 1 hr after mixing. **(E)** Quantification of fluorescence recovery of CsgF and CsgB after 1 hr incubation. Error bars shown are SD from 10 or more droplets. **(F)** Coomassie-stained SDS-PAGE of CsgF (20 μM) and CsgB (20 μM) alone or together. The SDS-PAGE wells of 30 min and 60 min CsgF (20 μM) +CsgB (20 μM) were highlighted in the red box.

## Discussion

Curli are bacterial functional amyloids found on the surface of enteric bacteria such as *E*.*coli* and *Salmonella spp* ^6^. Unlike disease-associated amyloids, curli production is well-orchestrated. CsgA protein, the major structural component of curli, is secreted to the cell surface in an unstructured state ^8^. CsgF is a curli accessory protein that assists the CsgB nucleator in providing a folding template that enables CsgA amyloid aggregation at the cell surface ^21^. Both CsgA and CsgB are secreted through the CsgG-CsgF outer membrane pore ^57^. NMR studies on CsgF have shown that the N-terminus and the C-terminus of CsgF are disordered (Figure 1B) ^47^. The conformational plasticity of IDRs makes them a better candidate for multiple functions ^58^. A large body of evidence has suggested that multivalent weak interactions involving proteins with disordered regions or low-complexity domains mediate phase separation ^39,50,52^. The presence of a prion-like low-complexity region in CsgF led us to hypothesize that CsgF might have the propensity to undergo phase separation (Figure 1C). CsgF readily formed phase separated droplets in a concentration dependent manner without any crowding agents (Figure 1D, 1E, and 1F). The N-terminus disordered region had a greater impact compared to the C-terminus in mediating CsgF phase separation (Figure 2B-E). CsgF function *in vivo* was affected by deletion of either the N-or C-terminus (Figure 3A-D). It has been shown for CsgA that its N-terminus mediates CsgA-CsgG interaction at the periplasm which allows translocation of CsgA to the cell surface ^23^. CsgA devoid of the N-terminal region (N-22 domain) is unstable in the periplasm ^57^. CsgF also secretes to the outer membrane via CsgG ^21^. The absence of CsgF-ΔN both in the whole cell and plug hinted that the CsgF N-terminus is essential for CsgG recognition and secretion. Thus, CsgF (without N-terminus) that are accumulated in the periplasm are susceptible to proteolytic degradation. Congo-red, exogenous complementation, and Western blot assays suggested that the N-terminal region is essential for the secretion of CsgF across the outer membrane, for the function of CsgF on the cell surface, and for the phase separation of CsgF.

Phase separation of prion-like domains is often described using a stickers-and-spacers model ^39,59,60^. Stickers are the amino acid residues that promote intra-and inter-molecular reversible networks, whereas spacers are the residues that are interspersed between stickers ^61^. Studies have shown that aromatic residues (Y/F) in the prion-like region act as stickers ^52,53^. The number and patterning of aromatic residues (valence) govern the phase behavior of prion-like domains ^52^. When phenylalanine residues in the N-terminus of CsgF were replaced with serine, a chemically neutral spacer, the phase separation activity of CsgF and curli biogenesis were both impaired (Figure 2G-J, 4, and S3B). A reduction in the propensity of CsgF to undergo phase separation with a decreasing number of phenylalanine residues indicated the importance of aromatic residues in modulating the phase-separation activity of CsgF (Figure 2G-J). The N-terminus of CsgF embeds on top of the CsgG opening at a 9:9 ratio ^22–24^, and single point mutations of phenylalanine residues at 24, 26, and 31 have shown to disrupt the CsgF interaction with CsgG ^23^. Our phenylalanine mutant proteins failed to complement *csgF* ^*-*^ cells when added exogenously (Figure 4A and 4B). The interactions of phenylalanine residues in the N-terminal region are essential for anchoring CsgF onto CsgG. The nonameric CsgG membrane channel could be a molecular crowder on the cell surface to bring CsgF molecules close enough to promote phase separation.

It was previously demonstrated that *csgA* ^*-*^ cells accept CsgA from donor cells (*csgB* ^*-*^ cells) in a process called interbacterial complementation ^12^. Therefore, we anticipated that *csgF* ^*-*^ cells might recruit CsgF to the cell surface if the protein is available outside the cell. Purified CsgF added exogenously complemented *csgF* ^*-*^ cells (Figure 3C-D). CsgF-ΔN or CsgF-ΔC failed to complement *csgF* ^*-*^ cells (Figure 3A-C). Thus, both the N- and C-termini are essential for the function of CsgF. Since the CsgF N-terminus is involved in CsgG interaction, it is likely that the C-terminus mediate interactions with CsgB. Immunofluorescence assays on curli expressing cells (MC4100) showed that CsgF localized on the cell surface as puncta, similar to what was observed previously in BW25113 (Figure 5F) ^24^. We find here that the presence of surface-localized CsgF regulates the secretion of CsgA to the outer membrane (Figure 5A-G).

CsgF is required for anchoring CsgB to the cell surface, without which cells fail to properly nucleate curli assembly ^12,21^. CsgF induces the formation of protease resistant CsgB on the cell surface ^21^. We find here that the curli nucleator protein CsgB does not phase separate on its own (Figure S4A). However, in the presence of CsgF, CsgB colocalized with CsgF condensates and transitioned from a dynamic state into a less dynamic amyloid state (Figure 6). Scaffolds promote assembly and maintain the integrity of biomolecular condensates, whereas clients are biomolecules that are recruited into condensates, but are not necessary for their formation ^37,62,63^. Clients with an affinity to scaffolds can be recruited into condensates ^62^. CsgB is not required to promote the phase separation of CsgF, but localized within the CsgF droplets (Figure 1D-E and 6A). Thus, CsgB is a client for the CsgF scaffold. It was shown previously using a pull-down assay that CsgF interacts with CsgB but not with CsgA ^23^. CsgA does not sequester to CsgF condensates like CsgB (Figure S4C). The CsgBΔR5 mutant (CsgB without C-terminus R5 repeat) of CsgB is also recruited to CsgF droplets (Figure S4B). In the absence of both the middle and C-terminal regions, CsgF does not interact with CsgB ^23^. Ectopic expression of ΔC-CsgF in a *csgF* ^*-*^ mutant strain revealed that ΔC-CsgF is bound to the cells, but does not complement *csgF* ^*-*^ cells to WT levels of curli production (Figure 3A-D), suggesting that the C-terminal 11-amino acid residues is interacting with the N-terminus of CsgB on the bacterial outer membrane. CsgB can nucleate the aggregation of CsgA ^8,9,55,64^. Based on this evidence, we propose that concentrating CsgB into CsgF droplets triggers the formation of an amyloid nuclei that promotes and spatially regulates CsgA aggregation specifically on the cell surface.

In this study, we demonstrate that the curli accessory protein CsgF undergoes phase separation. Phenylalanine residues in the N-terminal region are stickers, responsible for the assembly of CsgF condensates. Without the N-terminal region, CsgF failed to secrete to the cell surface, and both the N-terminal and C-terminal regions are required for the proper function of CsgF on the outer membrane. Exogenously added CsgF protein complemented *csgF* ^*-*^ cells. The curli nucleator protein CsgB acted as a client for the CsgF scaffold. CsgF condensates provided a platform to promote aggregation of CsgB (Figure 7). CsgF assembled on the outer membrane governed the secretion of CsgA. Taken together, this study provides insights into the interplay of multicomponent CsgF condensate and its influence on curli amyloid biogenesis.

**Figure 7.**
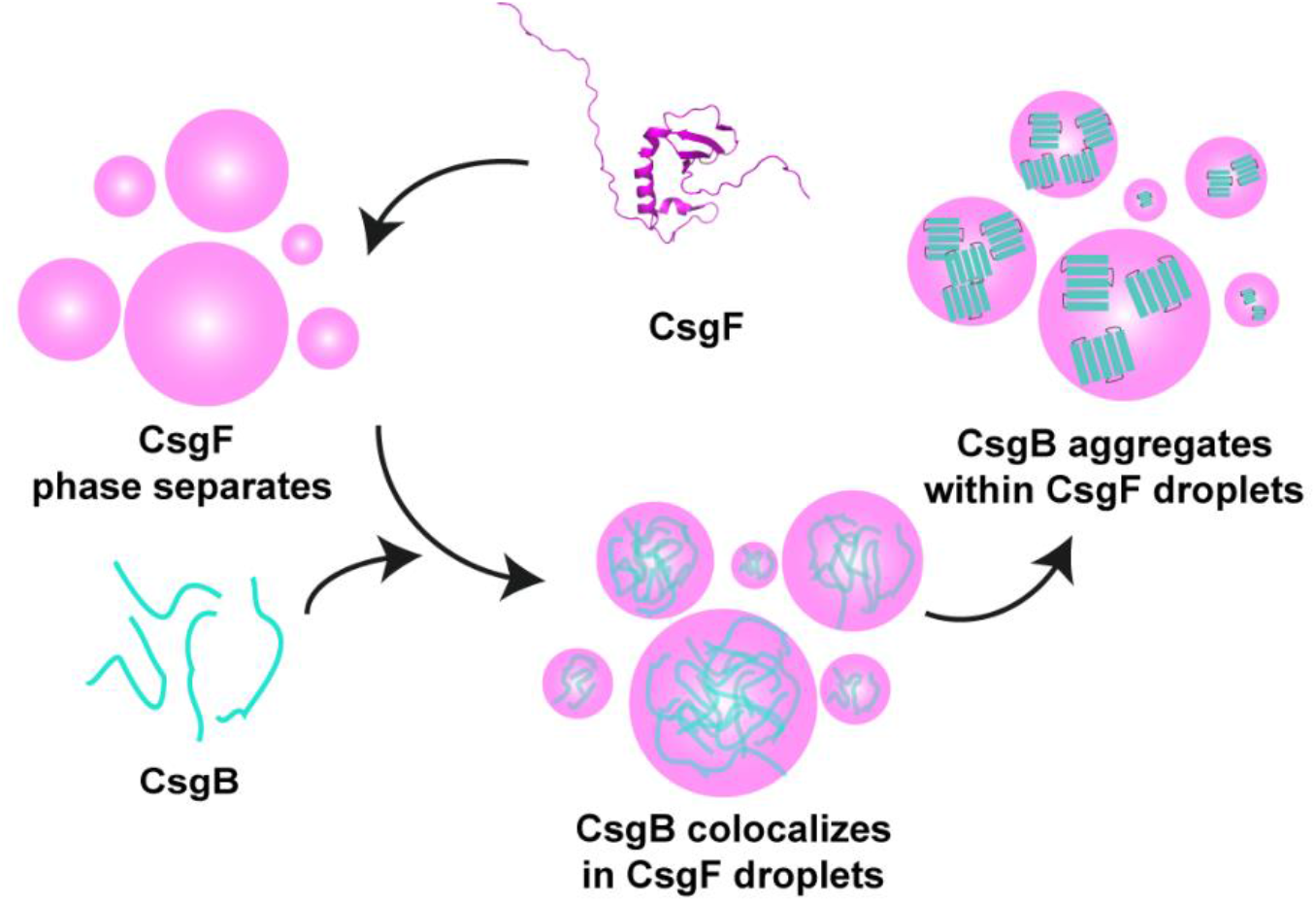
Model to show CsgB amyloid formation in the CsgF droplets. CsgF forms biocondensates. SDS-soluble monomeric CsgB is recruited in the CsgF droplets and undergoes amyloid formation.

## Materials and methods

### Protein purification

CsgF was expressed and purified as described previously with modifications ^47^. CsgF pET11d in BL21 (DE3) and CsgF variants were overexpressed using 500 μM IPTG for 4 hr at 37 ºC. The cell pellet from 250 mL culture was lysed in 25 mL of 8 M GdmCl, 150 mM NaCl, 50 mM potassium phosphate (KPi) pH 7.3, and the denatured solution was kept on a shaker overnight at RT. The lysate was centrifuged, and the supernatant was sonicated three times with pulse-on for 30 s and pulse-off 30 s. To the sonicated lysate 2.5 mL of Ni-NTA beads were added and allowed to bind for an hour at RT. Ni-NTA beads were washed with four different solutions. Wash 1: 10 mL of 8 M urea, 650 mM NaCl, 50 mM KPi pH 7.3. Wash 2: 6 mL of 4 M urea, 50 mM KPi pH 7.3. Wash 3: 6 mL of 2 M urea, 20 mM imidazole,50 mM KPi pH 7.3. Wash 4: 6 mL of 2 M urea, 40 mM imidazole,50 mM KPi pH 7.3. Then the protein was eluted in 2 M urea, 200 mM imidazole, 50 mM KPi pH 7.3. The eluents were dialyzed against 1% acetic acid and lyophilized. The protein was stored at -20 ºC for further use. The molar extinction coefficient used to estimate the concentration of CsgF, CsgF-ΔN, CsgF-ΔC were 3840 M^-1^cm^-1^, 2560 M^-1^cm^-1^, and 3840 M^-1^cm^-1^ (Scripps calculator), respectively. For all phenylalanine mutants, the molar extinction coefficient used was 3840 M^-1^cm^-1^. CsgA, CsgB, and CsgBΔR5 were purified as described elsewhere ^56,65^ and the concentrations were estimated by BCA assay.

### Labeling of proteins

Proteins were either dissolved or buffer exchanged in 8 M urea, 100 mM sodium carbonate pH 8.3 buffer, and kept at room temperature for 30 min with mild agitation. NHS-Alexa dyes or FITC (Invitrogen) to protein molar ratio used was 2:1. The reaction was carried out at room temperature for 1.5 hr. Free dye from the CsgF sample was removed by dialysis against 25 mM Tris pH 9 buffer. Dialyzed CsgF protein was then passed through a 5 mL Zeba spin desalting column (Thermo Fisher) that was equilibrated with 25 mM Tris pH 9 buffer. For CsgA and CsgB, 1 mL reaction mixture was diluted in 15 mL 8 M urea, 50 mM KPi pH 7.5 buffer, and concentrated using a 3 kDa filter (Sartorius) to remove excess dye. The concentrated solution was buffer exchanged with 50 mM KPi pH 7.3 buffer using a 5 mL Zeba spin desalting column. The concentration of labeled proteins was estimated using the molar extinction coefficient ε_494_ = 73,000 M^-1^cm^-1^ (Alexa-488), ε_632_ = 159,000 M^-1^cm^-1^ (Alexa-633), ε_280_ = 10810 M^-1^cm^-1^ (CsgA), ε_280_ = 7680 M^-1^cm^-1^(CsgB), and ε_280_ = 3840 M^-1^cm^-1^ (CsgB).

### Turbidity measurements

1.0 mM stocks of CsgF or CsgF variants were prepared in 1% acetic acid. The protein was diluted in 100 μL of 25 mM potassium phosphate pH 7.5 buffer and absorbance at 350 nm was recorded immediately using a TECAN infinite 200 PRO plate reader. Error bars shown are standard deviation from three independent measurements.

### Imaging and FRAP measurements

Imaging was performed on a 16-well glass-bottom culture well (Grace BioLabs) which was passivated with 5% (W/V) pluronic acid (Sigma Aldrich) for 2 hr and washed trice with 25 mM potassium phosphate pH 7.5 buffer. Labeled: Unlabeled protein used was 1:50. 1 mM CsgF was prepared in 1% acetic acid and diluted to the indicated buffer. Samples were imaged within 15-20 min either on a Leica DMI6000B inverted microscope (63x objective) or on a Nikon Ti2-E motorized inverted microscope (100x objective). For imaging CsgF together with CsgB, the required volume of CsgF was added from 100 μM CsgF, 25 mM potassium phosphate pH 7.5 (diluted from 1 mM CsgF, 1% acetic acid) to CsgB at room temperature. FRAP measurements were carried out using a Nikon Ti2-E motorized inverted microscope controlled by NIS Element software, a 100x objective (Oil CFI60 Plan Apochromat Lambda Series for DIC), a photometric Prime 95B Black-illuminated CMOS camera, and a SOLA 65 LED light source. Alexa-488 labeled CsgB was imaged using a GFP filter set [excitation, 470/40 nm (450 to 490 nm); emission, 525/50 nm (500 to 550 nm); dichroic mirror, 495 nm]. Alexa-633 labeled CsgF was imaged using a CY5 filter set [excitation, 620/60 nm (590 to 650 nm); emission, 700/75 nm (663 to 738 nm); dichroic mirror, 660 nm]. The region of interest was bleached with a 405-nm laser at 40% power (20 mW) with a 200-μs dwell time. Post-bleach images were acquired every 3 s for 5 min. The images were analyzed using ImageJ software. The bleached region was background corrected and normalized with the prebleached fluorescence intensity. The data was exported and plotted using OriginPro 2019 software. The error bars are the standard deviation from 10 or more bleached areas.

### Sedimentation assay

20 μM of 100 μL protein solution was prepared in 25 mM potassium phosphate pH 7.5 and allowed to stand at room temperature for 20 min. The protein samples were centrifuged at 16,0000 x g for 2 min. To the pellet, 100 μL of 8 M urea, 50 mM potassium phosphate pH 7.5 was added and the samples were boiled and subjected to electrophoresis on a 15% acrylamide gel. The Coomassie-stained gels was quantified using ImageJ.

### Congo red assay

The bacterial cells that were grown overnight with 50 μg/mL kanamycin were normalized to 1 OD_600_ in LB media and 4 μL cells were spotted on YESCA (<u>Y</u>east <u>E</u>xtract <u>C</u>asamino <u>A</u>cid) agar plate containing 50 μg/mL Congo red dye. For the exogenous complementation assay, protein stocks (1 μM, 10 μM, 20 μM, and 30 μM) were prepared in 25 mM Tris pH 9 buffer. 4 μL of protein was spotted on the YESCA-Congo red plate and air-dried. On the protein spot, 4 μL of 1 OD_600_ cells were spotted, and the plate was incubated at 26 ºC for 48 hr. The Congo red plates were imaged using a Canon EOS Rebel XSi camera and the background was changed using Adobe Photoshop 2022.

### Western blots and dot blots

The whole-cell and plug assay were carried out as described elsewhere ^65^. Briefly, the cells streaked on a YESCA agar plate and grown for 48 hr at 26 ºC were scraped off and resuspended in 50 mM potassium phosphate pH 7.3 buffer. 150 μL of cells (1 OD_600_) were centrifuged for 10 minutes at 16000 x g. To the cell pellet 100 μL of 88% formic acid was added and speed vacuumed to obtain formic acid treated whole cell samples. For the plug assay, the agar plate along with cells was stabbed with 1 mL pipette tip and carefully transferred to a 1.5 mL microcentrifuge tube using tweezers. 100 μL of 88% formic acid was added to the plug and vortexed until most of the plug was dissolved. The formic acid was removed from the samples using a speed vacuum at 55 ºC for 2 hr. To the formic acid and non-treated samples, 50 μL of 2x SDS-PAGE (polyacrylamide gel electrophoresis) running buffer and 10 μL of SDS-PAGE loading buffer were added. The samples were boiled for 10 minutes and subjected to electrophoresis. To the agar plugs that were treated with formic acid, 1 μL 0.5 M NaOH was added before boiling the samples. The transfer was carried out in 25 mM CAPS pH 11 buffer at 11 V for 15 hr. For CsgA antibody staining, PVDF membrane was used, and nitrocellulose membrane was used for probing CsgF. The dilutions used were 1:10,000 for CsgA antibody and 1:2500 for CsgF antibody. All the washes were performed in TBS-T buffer. For the intact cell dot blot, 4 μL of 1 OD_600_ cell were spotted on nitrocellulose membrane and blotting was performed in PBS buffer. IRDye 800CW goat anti-rabbit secondary antibody (Li-COR) at a 1:10,000 dilution was used, and the membranes were imaged using a Li-COR imaging system. The bands were quantified using ImageJ. Statistical analysis was performed using one way ANOVA with Tukey test in OriginPro 2019.

### Immunofluorescence staining

Cells were adhered to a polylysine-treated glass slide and fixed with 4% formaldehyde. The glass slides were blocked with 5% BSA for an hour and then treated with anti-6x His-tag antibody (1:100) for an hour at room temperature. After washing 3-times with PBS the cells were stained with the Alexa-568 labeled secondary antibody. The cells were washed thrice with PBS and stained with DAPI. Imaging was performed on a Leica SP8 inverted microscope with a 100x oil immersion objective lens.

## Acknowledgements

We would like to thank the current and the past members of the M.R.C. laboratory for helpful discussions and for strain and plasmid construction. We acknowledge Prof. Scott Hultgren (Washington University, St. Louis, MO) for providing us with the CsgF antibody. We would also like to thank Dr. Laura Buttitta (University of Michigan) for use of the Leica DMI6000 B microscope and Dr. Ajai Pulianmackal (Buttitta laboratory) for providing microscope training to H.M.S. We thank Prof. Samrat Mukhopadhyay (Indian Institute of Science Education and Research, Mohali, India) for critically reading the manuscript. This work is supported by NIH R01GM118651 and NIH grant R21AI137535 to M.R.C.

## Authors Contributions

H.M.S. and M.R.C. designed the project, J.L.B. collected FRAP measurements, C.E.D. purified and characterized some of the phenylalanine mutants, and H.M.S. performed all other experiments, M.R.C. acquired funding, A.G.V. and M.R.C. supervised, H.M.S. prepared figures and wrote the first draft, A.G.V. and M.R.C. edited the manuscript.

## References

1. Chiti, F. & Dobson, C. M. Protein Misfolding, Amyloid Formation, and Human Disease: A Summary of Progress Over the Last Decade. Annu. Rev. Biochem. 86, 27–68 (2017).

2. Hartl, F. U. Protein Misfolding Diseases. Annu. Rev. Biochem. 86, 21–26 (2017).

3. Hammer, N. D., Wang, X., McGuffie, B. A. & Chapman, M. R. Amyloids: Friend or Foe? J. Alzheimer’s Dis. 13, 407–419 (2008).

4. Balistreri, A., Goetzler, E. & Chapman, M. Functional Amyloids Are the Rule Rather Than the Exception in Cellular Biology. Microorganisms vol. 8 (2020).

5. Avni, A., Swasthi, H. M., Majumdar, A. & Mukhopadhyay, S. Chapter Five - Intrinsically disordered proteins in the formation of functional amyloids from bacteria to humans. In Dancing protein clouds: Intrinsically disordered proteins in health and disease, Part A (ed. Uversky, V. N. B. T.-P. in M. B.and T. S.) vol. 166 109–143 (Academic Press, 2019).

6. Barnhart, M. M. & Chapman, M. R. Curli Biogenesis and Function. Annu. Rev. Microbiol. 60, 131–147 (2006).

7. Kai-Larsen, Y. et al. Uropathogenic Escherichia coli Modulates Immune Responses and Its Curli Fimbriae Interact with the Antimicrobial Peptide LL-37. PLOS Pathog. 6, e1001010 (2010).

8. Bian, Z. & Normark, S. Nucleator function of CsgB for the assembly of adhesive surface organelles in Escherichia coli. EMBO J. 16, 5827–5836 (1997).

9. Hammer, N. D., Schmidt, J. C. & Chapman, M. R. The curli nucleator protein, CsgB, contains an amyloidogenic domain that directs CsgA polymerization. Proc. Natl. Acad. Sci. 104, 12494 LP – 12499 (2007).

10. Wang, X., Hammer, N. D. & Chapman, M. R. The Molecular Basis of Functional Bacterial Amyloid Polymerization and Nucleation *. J. Biol. Chem. 283, 21530–21539 (2008).

11. Hammar, M., Arnqvist, A., Bian, Z., Olsén, A. & Normark, S. Expression of two csg operons is required for production of fibronectin- and Congo red-binding curli polymers in Escherichia coli K-12. Mol. Microbiol. 18, 661–670 (1995).

12. Chapman, M. R. et al. Role of Escherichia coli curli operons in directing amyloid fiber formation. Science (80-.). 295, 851–855 (2002).

13. Bhoite, S., van Gerven, N., Chapman, M. R. & Remaut, H. Curli Biogenesis: Bacterial Amyloid Assembly by the Type VIII Secretion Pathway. EcoSal Plus 8, (2019).

14. Hiroshi, O., Kaneyoshi, Y. & Akira, I. Role of the Biofilm Master Regulator CsgD in Cross-Regulation between Biofilm Formation and Flagellar Synthesis. J. Bacteriol. 193, 2587–2597 (2011).

15. Loferer, H., Hammar, M. & Normark, S. Availability of the fibre subunit CsgA and the nucleator protein CsgB during assembly of fibronectin-binding curli is limited by the intracellular concentration of the novel lipoprotein CsgG. Mol. Microbiol. 26, 11–23 (1997).

16. Goyal, P. et al. Structural and mechanistic insights into the bacterial amyloid secretion channel CsgG. Nature 516, 250–253 (2014).

17. Epstein, E. A., Reizian, M. A. & Chapman, M. R. Spatial clustering of the curlin secretion lipoprotein requires curli fiber assembly. J. Bacteriol. 191, 608–615 (2009).

18. Klein, R. D. et al. Structure-function analysis of the curli accessory protein CsgE defines surfaces essential for coordinating amyloid fiber formation. MBio 9, (2018).

19. Evans, M. L. et al. The Bacterial Curli System Possesses a Potent and Selective Inhibitor of Amyloid Formation. Mol. Cell 57, 445–455 (2015).

20. Agarwal, A. & Mukhopadhyay, S. Prion Protein Biology Through the Lens of Liquid-Liquid Phase Separation. J. Mol. Biol. 434, 167368 (2022).

21. Nenninger, A. A., Robinson, L. S. & Hultgren, S. J. Localized and efficient curli nucleation requires the chaperone-like amyloid assembly protein CsgF. Proc. Natl. Acad. Sci. 106, 900 LP – 905 (2009).

22. Van der Verren, S. E. et al. A dual-constriction biological nanopore resolves homonucleotide sequences with high fidelity. Nat. Biotechnol. 38, 1415–1420 (2020).

23. Yan, Z., Yin, M., Chen, J. & Li, X. Assembly and substrate recognition of curli biogenesis system. Nat. Commun. 11, 241 (2020).

24. Zhang, M., Shi, H., Zhang, X., Zhang, X. & Huang, Y. Cryo-EM structure of the nonameric CsgG-CsgF complex and its implications for controlling curli biogenesis in Enterobacteriaceae. PLOS Biol. 18, e3000748 (2020).

25. Saier Jr., M. H. Microcompartments and Protein Machines in Prokaryotes. Microb. Physiol. 23, 243–269 (2013).

26. Azaldegui, C. A., Vecchiarelli, A. G. & Biteen, J. S. The emergence of phase separation as an organizing principle in bacteria. Biophys. J. 120, 1123–1138 (2021).

27. Cohan, M. C. & Pappu, R. V. Making the Case for Disordered Proteins and Biomolecular Condensates in Bacteria. Trends Biochem. Sci. 45, 668–680 (2020).

28. Yongdae, S. & P., B. C. Liquid phase condensation in cell physiology and disease. Science (80-.). 357, eaaf4382 (2017).

29. Hyman, A. A., Weber, C. A. & Jülicher, F. Liquid-Liquid Phase Separation in Biology. Annu. Rev. Cell Dev. Biol. 30, 39–58 (2014).

30. Boeynaems, S. et al. Protein Phase Separation: A New Phase in Cell Biology. Trends Cell Biol. 28, 420–435 (2018).

31. Bracha, D., Walls, M. T. & Brangwynne, C. P. Probing and engineering liquid-phase organelles. Nat. Biotechnol. 37, 1435–1445 (2019).

32. Lafontaine, D. L. J., Riback, J. A., Bascetin, R. & Brangwynne, C. P. The nucleolus as a multiphase liquid condensate. Nat. Rev. Mol. Cell Biol. 22, 165–182 (2021).

33. Mittag, T. & Pappu, R. V. A conceptual framework for understanding phase separation and addressing open questions and challenges. Mol. Cell 82, 2201–2214 (2022).

34. Kar, M. et al. Phase-separating RNA-binding proteins form heterogeneous distributions of clusters in subsaturated solutions. Proc. Natl. Acad. Sci. 119, e2202222119 (2022).

35. Guilhas, B. et al. ATP-Driven Separation of Liquid Phase Condensates in Bacteria. Mol. Cell 79, 293-303.e4 (2020).

36. Ladouceur, A.-M. et al. Clusters of bacterial RNA polymerase are biomolecular condensates that assemble through liquid–liquid phase separation. Proc. Natl. Acad. Sci. 117, 18540 LP – 18549 (2020).

37. Banani, S. F., Lee, H. O., Hyman, A. A. & Rosen, M. K. Biomolecular condensates: organizers of cellular biochemistry. Nat. Rev. Mol. Cell Biol. 18, 285–298 (2017).

38. Alberti, S., Gladfelter, A. & Mittag, T. Considerations and Challenges in Studying Liquid-Liquid Phase Separation and Biomolecular Condensates. Cell 176, 419–434 (2019).

39. Wang, J. et al. A Molecular Grammar Governing the Driving Forces for Phase Separation of Prion-like RNA Binding Proteins. Cell 174, 688-699.e16 (2018).

40. Martin, E. W. & Mittag, T. Relationship of Sequence and Phase Separation in Protein Low-Complexity Regions. Biochemistry 57, 2478–2487 (2018).

41. Martin, E. W. & Holehouse, A. S. Intrinsically disordered protein regions and phase separation: Sequence determinants of assembly or lack thereof. Emerg. Top. Life Sci. 4, 307–329 (2020).

42. Wang, B. et al. Liquid–liquid phase separation in human health and diseases. Signal Transduct. Target. Ther. 6, 290 (2021).

43. Lyon, A. S., Peeples, W. B. & Rosen, M. K. A framework for understanding the functions of biomolecular condensates across scales. Nat. Rev. Mol. Cell Biol. 22, 215–235 (2021).

44. Zbinden, A., Pérez-Berlanga, M., De Rossi, P. & Polymenidou, M. Phase Separation and Neurodegenerative Diseases: A Disturbance in the Force. Dev. Cell 55, 45–68 (2020).

45. de Oliveira, G. A. P., Cordeiro, Y., Silva, J. L. & Vieira, T. C. R. G. Chapter Nine - Liquid-liquid phase transitions and amyloid aggregation in proteins related to cancer and neurodegenerative diseases. In Protein Misfolding (ed. Donev, R. B. T.-A. in P. C. and S. B.) vol. 118 289–331 (Academic Press, 2019).

46. Ray, S. et al. α-Synuclein aggregation nucleates through liquid–liquid phase separation. Nat. Chem. 12, 705–716 (2020).

47. Schubeis, T. et al. Structural and functional characterization of the Curli adaptor protein CsgF. FEBS Lett. 592, 1020–1029 (2018).

48. Michelitsch, M. D. & Weissman, J. S. A census of glutamine/asparagine-rich regions: Implications for their conserved function and the prediction of novel prions. Proc. Natl. Acad. Sci. 97, 11910–11915 (2000).

49. Lancaster, A. K., Nutter-Upham, A., Lindquist, S. & King, O. D. PLAAC: a web and command-line application to identify proteins with prion-like amino acid composition. Bioinformatics 30, 2501–2502 (2014).

50. Shovamayee, M. et al. RNA buffers the phase separation behavior of prion-like RNA binding proteins. Science (80-.). 360, 918–921 (2018).

51. Krainer, G. et al. Reentrant liquid condensate phase of proteins is stabilized by hydrophobic and non-ionic interactions. Nat. Commun. 12, 1085 (2021).

52. Martin, E. W. et al. Valence and patterning of aromatic residues determine the phase behavior of prion-like domains. Science (80-.). 367, 694–699 (2020).

53. Holehouse, A. S., Ginell, G. M., Griffith, D. & Böke, E. Clustering of Aromatic Residues in Prion-like Domains Can Tune the Formation, State, and Organization of Biomolecular Condensates. Biochemistry 60, 3566–3581 (2021).

54. Yang, Y., Jones, H. B., Dao, T. P. & Castañeda, C. A. Single Amino Acid Substitutions in Stickers, but Not Spacers, Substantially Alter UBQLN2 Phase Transitions and Dense Phase Material Properties. J. Phys. Chem. B 123, 3618–3629 (2019).

55. Hammar, M., Bian, Z. & Normark, S. Nucleator-dependent intercellular assembly of adhesive curli organelles in Escherichia coli. Proc. Natl. Acad. Sci. 93, 6562 LP – 6566 (1996).

56. Swasthi, H. M. & Mukhopadhyay, S. Electrostatic lipid–protein interactions sequester the curli amyloid fold on the lipopolysaccharide membrane surface. J. Biol. Chem. 292, 19861–19872 (2017).

57. Robinson, L. S., Ashman, E. M., Hultgren, S. J. & Chapman, M. R. Secretion of curli fibre subunits is mediated by the outer membrane-localized CsgG protein. Mol. Microbiol. 59, 870–881 (2006).

58. Bondos, S. E., Dunker, A. K. & Uversky, V. N. On the roles of intrinsically disordered proteins and regions in cell communication and signaling. Cell Commun. Signal. 19, 88 (2021).

59. W., M. E. et al. Valence and patterning of aromatic residues determine the phase behavior of prion-like domains. Science (80-.). 367, 694–699 (2020).

60. Bremer, A. et al. Deciphering how naturally occurring sequence features impact the phase behaviours of disordered prion-like domains. Nat. Chem. 14, 196–207 (2022).

61. Choi, J.-M., Holehouse, A. S. & Pappu, R. V. Physical Principles Underlying the Complex Biology of Intracellular Phase Transitions. Annu. Rev. Biophys. 49, 107–133 (2020).

62. Posey, A. E., Holehouse, A. S. & Pappu, R. V. Chapter One - Phase Separation of Intrinsically Disordered Proteins. In Intrinsically Disordered Proteins (ed. Rhoades, E. B. T.-M.in E.) vol. 611 1–30 (Academic Press, 2018).

63. Ditlev, J. A., Case, L. B. & Rosen, M. K. Who’s In and Who’s Out—Compositional Control of Biomolecular Condensates. J. Mol. Biol. 430, 4666–4684 (2018).

64. Qin, S. et al. The E. coli CsgB nucleator of curli assembles to β-sheet oligomers that alter the CsgA fibrillization mechanism. Proc. Natl. Acad. Sci. 109, 6502–6507 (2012).

65. Zhou, Y., Smith, D. R., Hufnagel, D. A. & Chapman, M. R. Experimental manipulation of the microbial functional amyloid called curli. Methods Mol. Biol. 966, 53–75 (2013).

