## Supplementary material for "Cell Surface-localized CsgF Condensate is a Gatekeeper in Bacterial Curli Subunit Secretion": Supplemental Information.docx

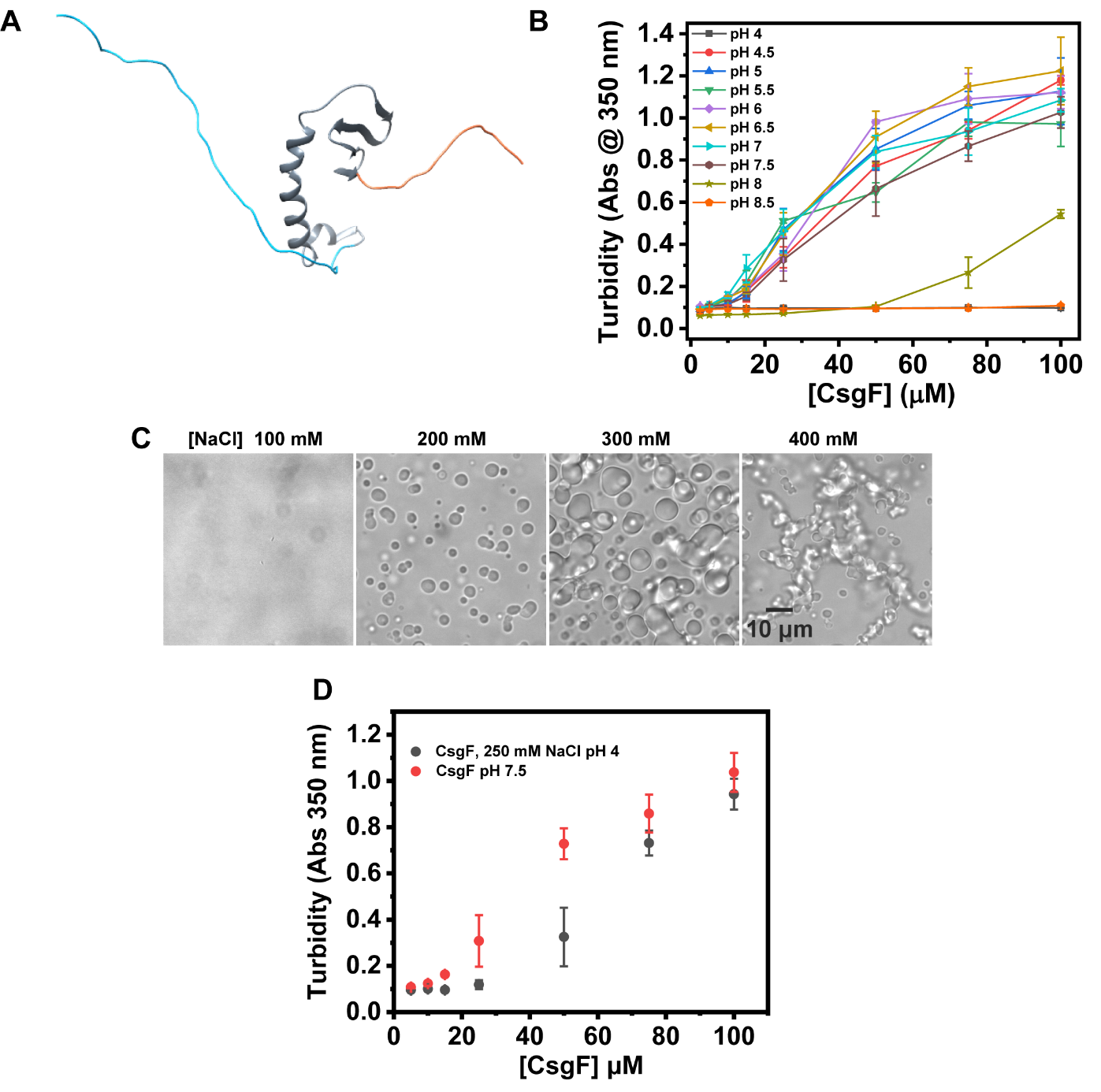

**Figure S1.** **(A)** The NMR structure of CsgF (PDB code: 5M1U). **(B)** Turbidity measurements of CsgF from pH 4 to 8.5. **(C)** DIC images of 100 µM CsgF at pH 4 in the presence of varying concentrations of NaCl. **(D)** Turbidity at varying concentrations of CsgF in 50 mM potassium phosphate pH 7.5 and in the presence of 250 mM NaCl, 25 mM sodium citrate pH 4.

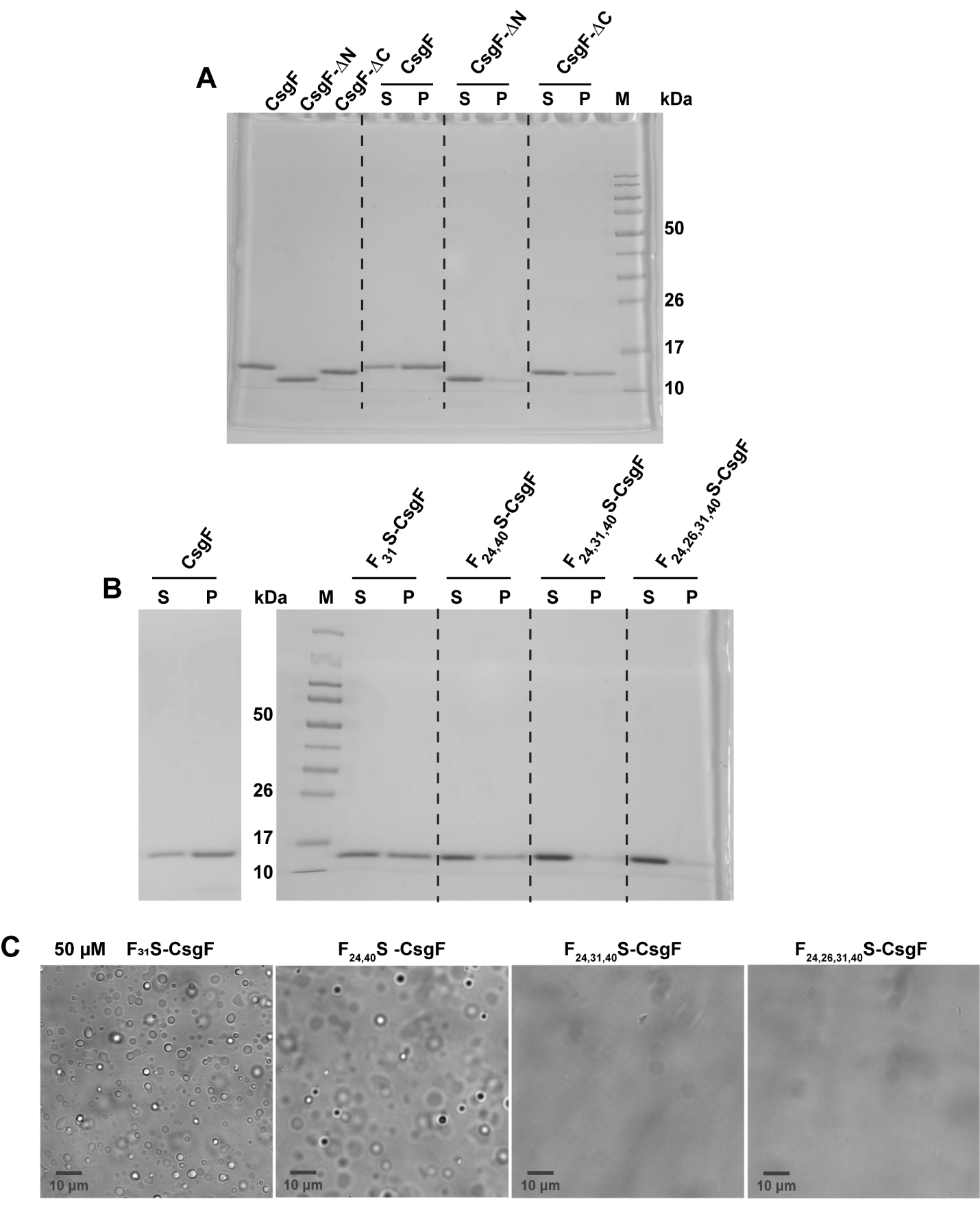

**Figure S2. (A)** Sedimentation assays performed on 20 µM CsgF, CsgF-∆N, and CsgF-∆C were run on SDS-PAGE and stained with Coomassie blue. **(B)** Coomassie-stained SDS-PAGE on the supernatant and pellet samples of 20 µM F_x_S-CsgF after sedimentation. **(C)** DIC images of 50 µM phenylalanine CsgF mutants in 50 mM potassium phosphate pH 7.5 buffer.

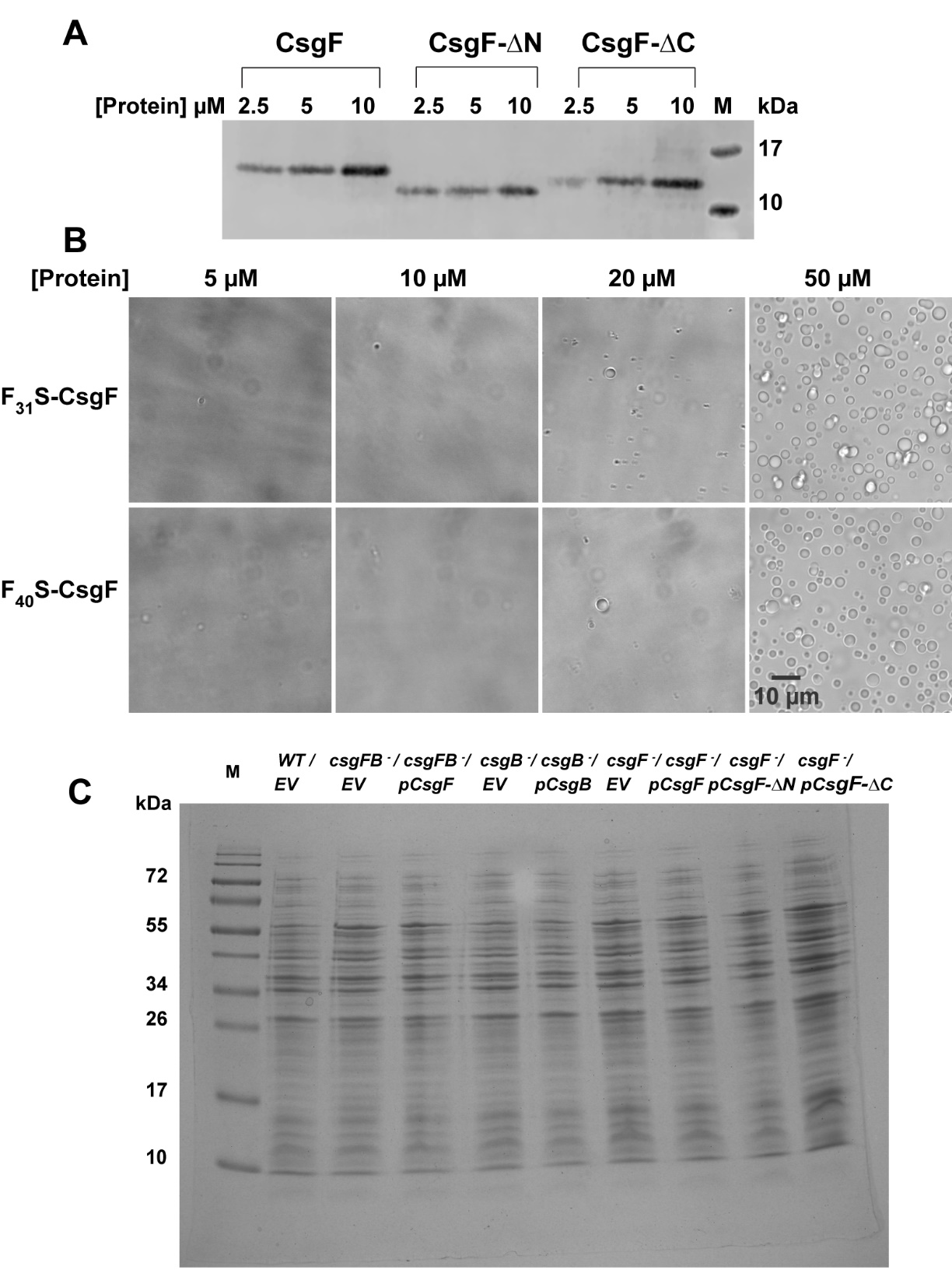

**Figure S3.** **(A)** Western blot on purified CsgF, CsgF-ΔN, and CsgF-ΔC probed with CsgF antibody on PVDF membrane. **(B)** DIC images of 5 µM, 10 µM, 20 µM, and 50 µM F31S-CsgF and F40S-CsgF **(C)** Coomassie-stained SDS-PAGE gel for the samples shown in Figure 5D.

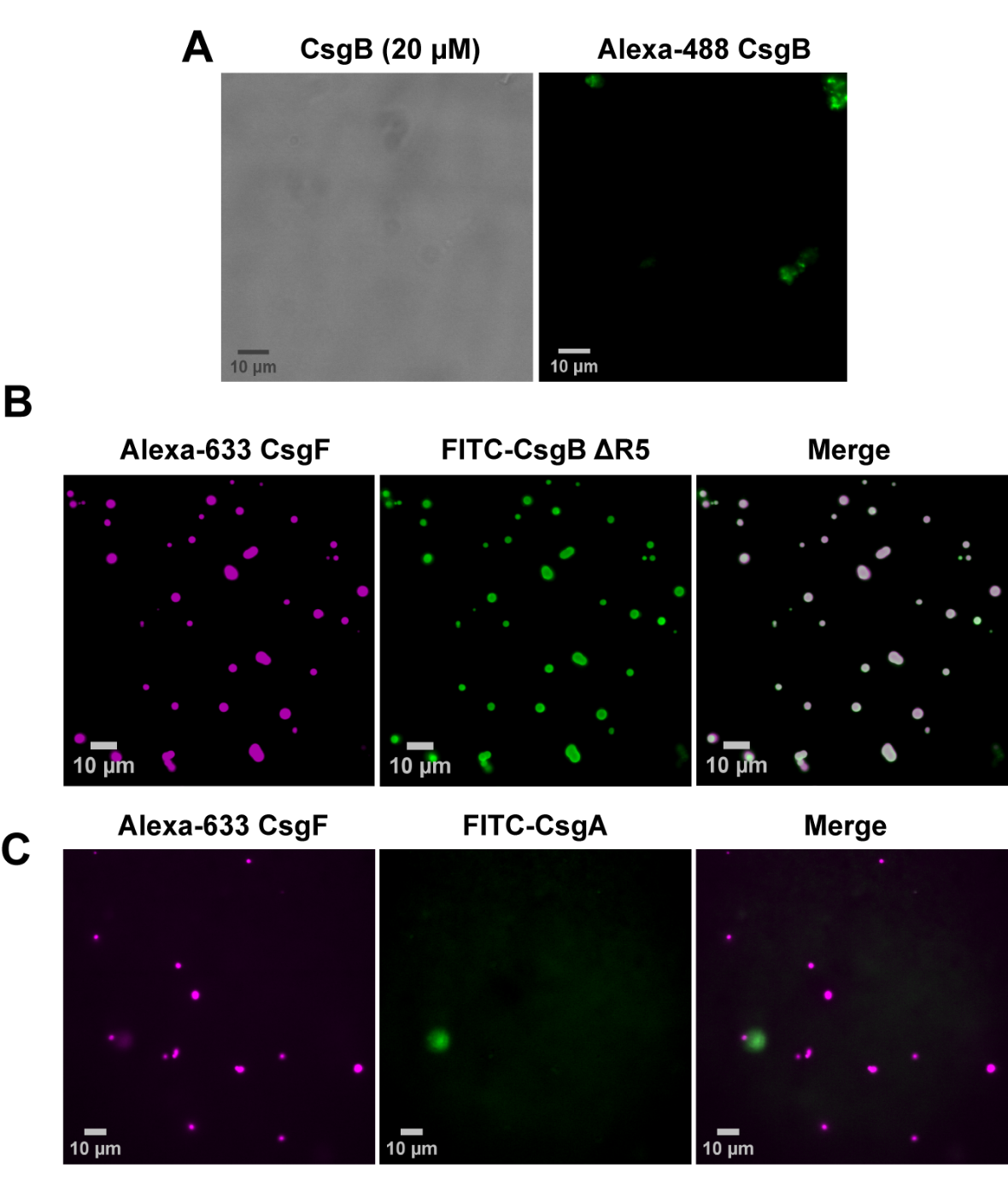

**Figure S4.** **(A)** DIC and fluorescence image of CsgB (20 µM) and Alexa-488 label CsgB. **(B)** Fluorescence images of CsgF (20 µM) and CsgB∆R5 (20 µM) (CsgB consist of 5 imperfect repeats and CsgB without the C-terminus R5 repeat region is named as CsgB ∆R5) **(C)** Fluorescence images of CsgF (20 µM) and CsgA (20 µM). The experiments were carried out in 50 mM potassium phosphate pH 7.5 buffer. The molar ratio of unlabeled to the labeled protein used was 50:1.

**Strains used in the study**

| **Common Name** | **Strains** | **Relevant Genotype** | **References** |
| --- | --- | --- | --- |
| Wild-type | MC4100 | F- *araD*139 Δ(*argF-lac*) U169 *rpsL*150(*strR*) *relA*1 *fibB*5301 *deoC*1 *ptsF*25 *rbsB* | ^1^ |
| *csgF^-^* | MHR592 | MC4100 *∆csgF* | ^2^ |
| *csgFB^-^* | MHR422 | MC4100 *∆csgF csgB* | ^2^ |
| *csgB^-^* | MHR261 | MC4100 *∆csgB* | ^3^ |
| BL21(DE3) |  | F-, ompT, hsdSB(rB-, mB-), dcm, gal, λ(DE3) | New England Biolabs |
| NEB3016 |  | MiniF *lacIq*(CamR) */ fhuA2 lacZ::T7 gene1 [lon] ompT gal sulA11 R(mcr-73::miniTn10--*TetS*)2 [dcm] R(zgb-210::Tn10--*TetS*) endA1 Δ(mcrC-mrr)114::IS10* | New England Biolabs |

**Plasmids used in the study**

| **Plasmids** | **Relevant characteristics** | **References** |
| --- | --- | --- |
| pET11d | IPTG-inducible expression vector | New England Biolabs |
| CsgF pET11d | C-terminal His­_6_ tagged *E.coli* CsgF cloned between Nco1/BamH1 sites of pET11d | ^4^ |
| CsgF-∆N pET11d | C-terminal His­_6_ tagged without 20 to 53 residues of CsgF cloned between Nco1/BamH1 sites of pET11d | This study |
| CsgF-∆C pET11d | C-terminal His­_6_ tagged without 127 to 138 residues of CsgF cloned between Nco1/BamH1 site of pET11d | This study |
| F_31_S CsgF pET11d | C-terminal His­_6_ tagged F31S CsgF cloned between Nco1/BamH1 site of pET11d | This study |
| F_24,40_S- CsgF pET11d | C-terminal His­_6_ tagged F24S and F40S CsgF cloned between Nco1/BamH1 site of pET11d | This study |
| F_24,31,40_S-CsgF pET11d | C-terminal His­_6_ tagged F24S, F31S, and F40S CsgF cloned between Nco1/BamH1 site of pET11d | This study |
| F_24,26,31,40_S-CsgF pET11d | C-terminal His­_6_ tagged F24S, F26S, F31S, and F40S CsgF cloned between Nco1/BamH1 site of pET11d | This study |
| CsgA pET11d | C-terminal His­_6_ tagged *E.coli* CsgA cloned between Nco1/BamH1 site of pET11d | ^5^ |
| CsgB pET11d | C-terminal His­_6_ tagged *E.coli* CsgB cloned between Nco1/BamH1 site of pET11d | ^6^ |
| CsgB ∆R5 | C-terminal His­_6_ tagged *E.coli* CsgB without R5 repeat was cloned between Nco1/BamH1 site of pET11d | ^7^ |
| pLR1 | *csgBA* promoter in pACYC177 vector | ^8^ |
| CsgF pLR1 | *E.coli* CsgF is cloned between Nco1/Pst1 of pLR1 | ^9^ |
| CsgF-∆N pLR1 | *E.coli* CsgF without 20 to 53 residues cloned between Nco1/Pst1 of pLR1 | This study |
| CsgF-∆C pLR1 | *E.coli* CsgF without 127 to 138 residues cloned between Nco1/Pst1 sites of pLR1 | This study |
| CsgB pLR2 | *E.coli* CsgF is cloned between Nco1/BamH1 of pLR2 | ^7^ |
| pTrc99A | IPTG inducible expression vector | Pharmacia Biotech |
| CsgF-His pTrc99A | E.coli Csg-His cloned between Ecor1 and BamH1 sites of pTrc99A | ^7^ |

**Primers used in the study**

| **Primers** | **Primer sequence** |
| --- | --- |
| His ΔN-CsgF F  His ΔN-CsgF R | 5’ CATGCCATGGCAAGCTATAACGATGAC 3’  5’ GCCGGATCCTTAGTGATGGTGATG 3’ |
| His ΔC-CsgF F  His ΔC-CsgF R | 5’ TCGACCATCCAGCATCACCATCACCATC 3’  5’ TGATGGTGATGCTGGATGGTCGAGGTTTG 3’ |
| CsgF F24S F  CsgF F24S R | 5’ GAACCATGACTAGCCAGTTCCGTAATC 3’  5’ GATTACGGAACTGGCTAGTCATGGTTC 3’ |
| CsgF F26S F  csgF F26S R | 5’ GACTTTCCAGAGCCGTAATCCAAAC 3’  5’ GTTTGGATTACGGCTCTGGAAAGTC 3’ |
| CsgF F31S F  CsgF F31S R | 5’ GTAATCCAAACAGTGGTGGTAAC 3’  5’ GTTACCACCACTGTTTGGATTAC 3’ |
| CsgF F40S F  CsgF F40S R | 5’ CCAAATAATGGCGCTAGTTTATTAAATAGC 3’  5’ GCTATTTAATAAACTAGCGCCATTATTTGG 3’ |
| Sec-CsgF-∆N F  Sec-CsgF-∆N R | 5’ GTTGGGCTAGCTATAACGATGACTTTGG 3’  CATCGTTATAGCTAGCCCAACTTAATGG |
| Sec-CsgF-∆C F  Sec-CsgF-∆C R | 5’ TATTGACGACGGGATCAGTACC 3’  5’ CAAACTGCAGTTACTGGATGGTCGAGGTTTGTCC 3’ |

**References**

1.	Casadaban, M. J. Transposition and fusion of the lac genes to selected promoters in Escherichia coli using bacteriophage lambda and Mu. *J. Mol. Biol.* **104**, 541–555 (1976).

2.	R., C. M. *et al.* Role of Escherichia coli Curli Operons in Directing Amyloid Fiber Formation. *Science (80-. ).* **295**, 851–855 (2002).

3.	Hammar, M., Arnqvist, A., Bian, Z., Olsén, A. & Normark, S. Expression of two csg operons is required for production of fibronectin- and Congo red-binding curli polymers in Escherichia coli K-12. *Mol. Microbiol.* **18**, 661–670 (1995).

4.	Schubeis, T. *et al.* Structural and functional characterization of the Curli adaptor protein CsgF. *FEBS Lett.* **592**, 1020–1029 (2018).

5.	Cegelski, L. *et al.* Small-molecule inhibitors target Escherichia coli amyloid biogenesis and biofilm formation. *Nat. Chem. Biol.* **5**, 913–919 (2009).

6.	Qin, S. *et al.* The E. coli CsgB nucleator of curli assembles to β-sheet oligomers that alter the CsgA fibrillization mechanism. *Proc. Natl. Acad. Sci.* **109**, 6502–6507 (2012).

7.	Hammer, N. D., Schmidt, J. C. & Chapman, M. R. The curli nucleator protein, CsgB, contains an amyloidogenic domain that directs CsgA polymerization. *Proc. Natl. Acad. Sci.* **104**, 12494 LP – 12499 (2007).

8.	Robinson, L. S., Ashman, E. M., Hultgren, S. J. & Chapman, M. R. Secretion of curli fibre subunits is mediated by the outer membrane-localized CsgG protein. *Mol. Microbiol.* **59**, 870–881 (2006).

9.	A., N. A., S., R. L. & J., H. S. Localized and efficient curli nucleation requires the chaperone-like amyloid assembly protein CsgF. *Proc. Natl. Acad. Sci.* **106**, 900–905 (2009).
